# Deciphering the Mechanistic Continuum of Broadly Neutralizing Class 4 Antibodies Targeting Conserved Cryptic Epitopes of the SARS-CoV-2 Spike Protein : Operating at the Intersection of Binding, Allostery and Immune Escape

**DOI:** 10.1101/2025.07.10.664240

**Authors:** Mohammed Alshahrani, Vedant Parikh, Brandon Foley, Gennady Verkhivker

**Affiliations:** Keck Center for Science and Engineering, Graduate Program in Computational and Data Sciences, Schmid College of Science and Technology, Chapman University, Orange, CA 92866, United States of America; Department of Biomedical and Pharmaceutical Sciences, Chapman University School of Pharmacy, Irvine, CA 92618, United States of America

## Abstract

Understanding atomistic basis of multi-layer mechanisms employed by broadly reactive neutralizing antibodies of SARS-CoV-2 spike protein without directly blocking receptor engagement remains an important challenge in coronavirus immunology. Class 4 antibodies represent an intriguing case: they target a deeply conserved, cryptic epitope on the receptor-binding domain yet exhibit variable neutralization potency across subgroups F1 (CR3022, EY6A, COVA1-16), F2 (DH1047), and F3 (S2X259). In this study, we employed a multi-modal computational approach combining atomistic and coarse-grained simulations, mutational scanning of binding interfaces, binding free energy calculations, and allosteric modeling using dynamic network analysis to map the allosteric landscapes and binding hotspots of these antibodies. Through this approach, our data revealed that distal binding can influence ACE2 engagement and immune escape traits through the confluence of direct interfacial interactions and allosteric effects. We found that group F1 antibodies can operate via classic allostery by modulating flexibility in the receptor binding domain loop regions and indirectly interfering with ACE2 binding using long-range effects. Group F2 antibody DH1047 represents an intermediate mechanism, engaging residues T376, R408, V503, and Y508 hotspots which are both critical for ACE2 binding and under immune pressure. Mutational scanning and rigorous binding free energy calculations highlight the synergy between hydrophobic and electrostatic interactions, while dynamic network modeling reveals a shift toward localized communication pathways connecting the cryptic site to the ACE2 interface. Our results demonstrate how group F3 antibody S2X259 achieves efficient synergistic mechanism through confluence of direct competition with ACE2 and localized allosteric effects leading to stabilization of the spike protein at the cost of increased escape vulnerability. Dynamic network analysis identifies a shared “allosteric ring” embedded in the core of the receptor binding domain and serving a conserved structural scaffold mediating long-range signal propagation with antibody-specific extensions propagating toward the ACE2 interface. The findings of this study support a modular model of antibody-induced allostery where neutralization strategies evolve via refinement of peripheral network connections, rather than complete redesign of the epitope itself. Taken together, this study establishes a robust computational framework for atomistic understanding of mechanisms underlying neutralization activity and immune escape for class 4 antibodies which harness conformational dynamics, binding energetics, and allosteric modulation to influence viral entry. The findings highlight the modular evolution of neutralization strategies, where progressive refinement of peripheral interactions enhances potency but increases susceptibility to immune pressure.

## Introduction

The emergence of severe acute respiratory syndrome coronavirus 2 (SARS-CoV-2) has triggered an unprecedented surge in global research efforts aimed at deciphering its molecular architecture, mechanisms of cellular invasion, and the immune responses it elicits. Central to these investigations is the SARS-CoV-2 Spike (S) glycoprotein, a trimeric surface protein that plays a pivotal role in mediating viral entry into host cells and serves as the primary target for neutralizing antibodies [1–15]. The S protein exhibits extraordinary conformational plasticity, enabling it to navigate through multiple functional states—from receptor engagement to membrane fusion—while simultaneously evading immune surveillance. Structurally, the S protein is composed of two functionally distinct subunits: S1 and S2. The S1 subunit contains four key domains—the N-terminal domain (NTD), the receptor-binding domain (RBD), and two conserved subdomains, SD1 and SD2 —each contributing uniquely to the dynamic life cycle of the virus. The NTD is implicated in early stages of host cell recognition, potentially facilitating attachment via interactions with glycans or other cell surface components [1–15]. However, the RBD remains the focal point of both infection and immunity, as it undergoes reversible transitions between “up” and “down” conformations. In the “up” state, the RBD becomes exposed and accessible for binding to the angiotensin-converting enzyme 2 (ACE2) receptor on host cells, while in the “down” state, it is shielded from antibody recognition [1–15]. Beyond the RBD, the SD1 and SD2 subdomains play essential structural roles in maintaining the prefusion conformation of the S protein, acting as molecular scaffolds that regulate the timing and efficiency of membrane fusion [10–18]. Detailed biophysical analyses have illuminated the thermodynamic and kinetic underpinnings of these conformational transitions, revealing how subtle energetic shifts govern large-scale structural rearrangements critical for viral infectivity [16–18]. Over the course of the pandemic, the rapid evolution of SARS-CoV-2 variants of concern (VOCs) has underscored the importance of understanding how mutations in the S protein affect antigenicity and transmissibility. Cryo-electron microscopy (cryo-EM) and X-ray crystallography studies generated an extensive structural atlas of the S protein in various functional states, including complexes formed with neutralizing antibodies [19–25]. A cornerstone of the adaptive immune response to SARS-CoV-2 is the production of neutralizing antibodies targeting the S protein, especially those directed against the RBD. Recent technological innovations such as high-throughput yeast display screening and deep mutational scanning (DMS) have revolutionized our ability to dissect the antigenic landscape of the RBD at near-residue resolution [26]. Through the identification of 247 broadly neutralizing antibodies against sarbecoviruses. combined with high-throughput yeast-display DMS the escape mutation profiles of the RBD were determined and these antibodies into six major epitope groups (A–F) based on their breadth and epitope specificity [26]. This classification was revisited using data from post-vaccination infections with the Omicron BA.1 variant, leading to the identification of 12 distinct epitope groups among 1,640 RBD-targeting antibodies [27]. This expanded categorization provides a more nuanced understanding of how different antibody classes interact with the RBD and respond to emerging mutations. Among these groups, A–C are characterized by antibodies that bind directly within the ACE2-binding motif, making them highly effective at blocking viral attachment. However, they are also the most vulnerable to escape mutations, particularly at residues K417, E484, and N501, which are frequently altered in VOCs such as Beta, Gamma, and Omicron lineages. These mutations can significantly reduce antibody binding affinity and neutralization potency, highlighting the evolutionary pressure exerted by immune selection. In contrast, Group D antibodies, exemplified by monoclonals such as REGN-10987, LY-CoV1404, and COV2-2130, recognize a distinct epitope centered around residues 440–449 on the RBD. This group is further subdivided into D1 and D2 subgroups, reflecting subtle differences in binding orientation and mutational sensitivity. Groups E and F represent antibodies that recognize epitopes outside the central ACE2-binding site, offering broader protection due to their less direct overlap with the receptor interface. These groups were further subdivided into E1–E3 and F1–F3, respectively, covering both the front and back surfaces of the RBD, thereby providing a comprehensive view of non-competitive epitopes. In this classification, group E1 antibodies target the S309 epitope [28,29], group E3 antibodies target the S2H97 binding site [30], group F1 antibodies recognize the CR3022/S304 epitope [31,32], group F2 antibodies bind to the DH1047 site [33], and group F3 antibodies binding the ADG-2 site [34,35]. The epitope clusters E and F correspond to Class 3 and Class 4 antibodies in earlier taxonomies [36]. The follow-up seminal studies analyzed the escape mutation profiles of a total of 2,688 monoclonal antibodies including 1,874 antibodies isolated from individuals infected with XBB or JN.1 variants, ultimately resulting in the identification of 22 distinct antibody clusters and a detailed antigenic map of the RBD [37,38]. Importantly, this latest pioneering study showed the possibility of accurately predicting SARS-CoV-2 RBD evolution by aggregating high-throughput antibody DMS results and constructing pseudoviruses that carry the predicted mutations as filters to screen for potent neutralizing antibodies. Subsequent discoveries of E1 and F3 antibodies [39,40] revealed that group F3 antibody SA55 and group E1 antibody SA58 can bind non-competitively to the RBD with robust neutralizing activity against a broad range of immune-evading variants, including those harboring convergent mutations in the RBD of BQ.1, BQ.1.1, and XBB lineages [39,40]. Another large-scale investigation employing high-throughput DMS assays for 1,637 potent monoclonal antibodies evaluated immune escape across eight major SARS-CoV-2 variants, including B.1 (D614G), Omicron BA.1, BA.2, BA.5, BQ.1.1, XBB.1.5, HK.3, and JN.1 [41]. Pan-sarbecovirus binding assays combined with in vitro mapping of viral escape, structural analyses and DMS experiments provided a comprehensive characterization of a panel of antibodies targeting different epitopes, including conserved cryptic RBD region [42]. In a recent breakthrough, two newly identified antibodies CYFN1006-1 and CYFN1006-2 demonstrated superior neutralization breadth across all tested SARS-CoV-2 variants, even outperforming SA55 [43]. Their distinct binding profile suggests that combining SA55 with CYFN1006-1 could offer synergistic protection against JN.1, KP.2, KP.3, and other emerging SARS-CoV-2 variants [43]. These investigations have systematically cataloged RBD escape mutations associated with a wide array of monoclonal antibodies, revealing distinct patterns of escape-prone regions and mutation-resistant epitopes. Recent surveillance of emerging variants indicates that SARS-CoV-2 preferentially accumulates mutations in frequently targeted “hotspot” epitopes, while maintaining the structural integrity of critical RBD regions essential for folding, stability, and function.

Computational approaches have played a transformative role in elucidating the structural dynamics and functional mechanisms of the SARS-CoV-2 S protein at atomic resolution. These tools have enabled researchers to probe not only the intrinsic flexibility of the S protein but also its interactions with key molecular partners such as the ACE2 receptor and a wide array of neutralizing antibodies. Through advanced simulation techniques—including molecular dynamics (MD) simulations and Markov state models (MSMs) —the conformational landscapes of Omicron subvariants such as XBB.1 and XBB.1.5, both in their free forms and bound to ACE2 or antibodies, have been systematically characterized [44]. These studies have provided critical insights into how mutations modulate conformational transitions, stability, and receptor accessibility across evolving variants. In parallel, computational mutational scanning and binding affinity analyses have offered a quantitative framework for interpreting experimental observations related to the interaction between Omicron XBB variants and ACE2, as well as their susceptibility to a panel of Class 1 neutralizing antibodies [45,46]. These approaches have proven instrumental in dissecting the molecular basis of immune escape and viral adaptation. Building on these advances, our group has integrated AlphaFold2-based atomistic modeling with ensemble sampling methods to predict and analyze the structures and conformational ensembles of S-ACE2 complexes across the most prevalent Omicron subvariants, including JN.1, KP.1, KP.2, and KP.3 [47]. Recent investigations uncovered a multifactor-based mechanism that governs the emergence of escape mutants against ultra-potent monoclonal antibodies [48,49].In the proposed mechanism, the selection of specific mutations is driven by an intricate interplay between their effects on protein structural stability, binding affinity, and long-range allosteric communication networks within the RBD. Further computational efforts have focused on understanding the mechanisms of broadly neutralizing antibodies, particularly those belonging to the E1 and F3 groups [50]. A recent comparative modeling study shed light on the distinct molecular strategies employed by several broadly neutralizing antibodies, including S309, S304, CYFN1006, and VIR-7229 [51]. These findings revealed two overarching paradigms: conservation-driven binding, wherein antibodies target highly conserved residues crucial for viral function, and adaptability-driven binding, where antibodies exploit flexible or dynamic interfaces to maintain potency despite antigenic drift. Computational and experimental studies highlighted many factors underlying the complexity of SARS-CoV-2 evolution at the molecular level [52–54]. Conformational dynamics and allosteric perturbations can be linked to binding of novel human antibodies where antibody-induced dynamics can render weak, moderate and strong neutralizing antibodies [55]. Collectively, the existing evidence suggests that viral evolution is not merely a stochastic process but rather a finely tuned balance between immune evasion, receptor-binding affinity, and fitness constraints imposed by mutation-induced structural perturbations. These forces are further shaped by the diversity and specificity of the host antibody repertoire.

In the current study, we investigate the interplay between dynamic, energetic, and allosteric mechanisms that govern antibody binding and immune escape, focusing specifically on distinct groups of Class 4 antibodies. Class 4 antibodies represent a unique category of broadly reactive monoclonal antibodies that target a deeply conserved, cryptic epitope within the S-RBD protein. Here, we present a multi-dimensional analysis integrating structural mapping, conformational dynamics, mutational scanning, and binding free energy calculations, along with dynamic network modeling, to elucidate the molecular basis of antibody recognition, allostery, and immune escape across class 4 group F1 (CR3022, EY6A, COVA1-16), group F2 (DH1047), and group F3 (S2X259) antibodies. We perform structural binding epitope mapping and analysis of closely-related cross-reactive but weakly neutralizing class 4 group F1 antibodies CR3022 [31,56,57], EY6A [57] and COVA1-16 [58]. Through structural analysis we establish that in addition to CR3022, EY6A and COVA1-16 class 4 antibodies targeting the conserved cryptic site with very similar binding footprints can be all classified as group F1 [ 26,27,37–39]. Additionally, we studied class 4 antibodies DH1047 of group F2 [33], and S2X259 of group F3 [59].The classification of RBD-directed antibodies has recently been redefined to include a larger set of antibodies and finer epitope binning [60]. To investigate dynamics, energetics and binding profiles of these antibodies with S-RBD we employed coarse-grained (CG) and atomistic molecular dynamics (MD) simulations, systematic mutational scanning of the antibody-RBD binding interfaces and binding free energy calculations. Through network-based modeling of conformational ensembles, we explored the allosteric consequences of antibody binding. Our results provide a broader mechanistic framework for understanding how binding and allostery synergistically shape immune defense responses, which are inherently context-dependent and energetically nuanced. This work establishes a mechanistic continuum among class 4 antibodies, from indirect allostery (F1) to hybrid (F2) and combination of direct receptor blocking with localized allostery (F3), revealing key insights into how cryptic site recognition translates into functional impact. Importantly, it highlights that neutralization need not rely solely on physical occlusion of ACE2 but can emerge through dynamic reorganization of the RBD. These findings offer a useful computational framework for understanding antibody function beyond canonical epitopes and provide guidance for designing next-generation therapeutics and vaccines that harness the power of conformational dynamics and network-level control to combat evolving sarbecoviruses.

## Materials and Methods

### Coarse-Grained Simulations Molecular Dynamics Simulations

The crystal and cryo-EM structures of the RBD-antibody are obtained from the Protein Data Bank [61] We employed CABS-flex approach that efficiently combines a high-resolution coarse-grained model and efficient search protocol capable of accurately reproducing all-atom MD simulation trajectories and dynamic profiles of large biomolecules on a long time scale [62–67]. In this high-resolution model, the amino acid residues are represented by Cα, Cβ, the center of mass of side chains and another pseudoatom placed in the center of the Cα-Cα pseudo-bond. In this model, the amino acid residues are represented by Cα, Cβ, the center of mass of side chains and the center of the Cα-Cα pseudo-bond. The CABS-flex approach implemented as a Python 2.7 object-oriented standalone package was used in this study to allow for robust conformational sampling proven to accurately recapitulate all-atom MD simulation trajectories of proteins on a long time scale. Conformational sampling in the CABS-flex approach is conducted with the aid of Monte Carlo replica-exchange dynamics and involves local moves of individual amino acids in the protein structure and global moves of small fragments. The default settings were used in which soft native-like restraints are imposed only on pairs of residues fulfilling the following conditions : the distance between their *C*_α_ atoms was smaller than 8 Å, and both residues belong to the same secondary structure elements. A total of 1000 independent CG-CABS simulations were performed for each of the systems studied. In each simulation, the total number of cycles was set to 10,000 and the number of cycles between trajectory frames was 100. MODELLER-based reconstruction of simulation trajectories to all-atom representation [68] provided by the CABS-flex package was employed to produce atomistic models of equilibrium ensembles for studied systems.

### Molecular Dynamics Simulations

The missing regions for the studied structures of the RBD-antibody are reconstructed and optimized using template-based loop prediction approach ArchPRED [69]. The side chain rotamers were refined and optimized by SCWRL4 tool [70]. The protonation states for all the titratable residues of the antibody and RBD proteins were predicted at pH 7.0 using Propka 3.1 software and web server [71,72]. The glycan chains were built using CHARMM-GUI Glycan Reader [73,74] and Modeller [68] at glycosylation sites N331 and N343 of RBD. NAMD 2.13-multicore-CUDA package [75] with CHARMM36m force field [76] employed to perform all-atom MD simulations for the RBD-antibody complexes. Each system was solvated with TIP3P water molecules and neutralizing 0.15 M NaCl in a periodic box that extended 10 Å beyond any protein atom in the system [77]. All Na+ and Cl− ions were placed at least 8 Å away from any protein atoms and from each other. MD simulations are typically performed in an aqueous environment in which the number of ions remains fixed for the duration of the simulation, with a minimally neutralizing ion environment or salt pairs to match the macroscopic salt concentration [78]. The heavy atoms in the complex were restrained using a force constant of 1000 kJ mol^−1^ nm^−1^ to perform 500 ps equilibration simulation. Long-range, non-bonded van der Waals interactions were computed using an atom-based cutoff of 12 Å, with the switching function beginning at 10 Å and reaching zero at 14 Å. The SHAKE method was used to constrain all the bonds associated with hydrogen atoms. The simulations were run using a leap-frog integrator with a 2 fs integration time step. The ShakeH algorithm in NAMD was applied for the water molecule constraints. A 310 K temperature was maintained using the Nóse-Hoover thermostat with 1.0 ps time constant and 1 atm pressure was maintained using isotropic coupling to the Parrinello-Rahman barostat with time constant of 5.0 ps [79,80]. The long-range electrostatic interactions were calculated using the particle mesh Ewald method [81] with a cut-off of 1.2 nm and a fourth-order (cubic) interpolation. The simulations were performed under an NPT ensemble with a Langevin thermostat and a Nosé–Hoover Langevin piston at 310 K and 1 atm. The damping coefficient (gamma) of the Langevin thermostat was 1/ps. In NAMD, the Nosé–Hoover Langevin piston method is a combination of the Nosé– Hoover constant pressure method [82] and piston fluctuation control implemented using Langevin dynamics [83]. An NPT production simulation was run on equilibrated structures for 1µs keeping the temperature at 310 K and a constant pressure (1 atm).

### Mutational Scanning Profiling

We conducted mutational scanning analysis of the binding epitope residues for the S RBD-antibody complexes. Each binding epitope residue was systematically mutated using all substitutions and corresponding protein stability and binding free energy changes were computed. BeAtMuSiC approach [84–88] was employed that is based on statistical potentials describing the pairwise inter-residue distances, backbone torsion angles and solvent accessibilities, and considers the effect of the mutation on the strength of the interactions at the interface and on the overall stability of the complex. The BeAtMuSiC approach evaluates the impact of mutations on both the strength of interactions at the protein-protein interface and the overall stability of the complex using statistical energy functions for ΔΔ*G* estimation, derived from the Boltzmann law which relates the frequency of occurrence of a structural pattern to its free energy. BeAtMuSiC identifies a residue as part of the protein-protein interface if its solvent accessibility in the complex is at least 5% lower than its solvent accessibility in the individual protein partner(s). The binding free energy of protein-protein complex can be expressed as the difference in the folding free energy of the complex and folding free energies of the two protein binding partners:

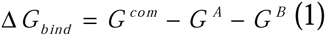

The change of the binding energy due to a mutation was calculated then as the following:

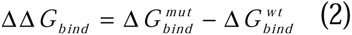

We leveraged rapid calculations based on statistical potentials to compute the ensemble-averaged binding free energy changes using equilibrium samples from simulation trajectories. The binding free energy changes were obtained by averaging the results over 1,000 and 10, 000 equilibrium samples for each of the systems studied.

### Binding Free Energy Computations

We calculated the ensemble-averaged changes in binding free energy using 1,000 equilibrium samples obtained from simulation trajectories for each system under study. Initially, the binding free energies of the RBD-antibody complexes were assessed using the MM-GBSA approach [89,90]. Additionally, we conducted an energy decomposition analysis to evaluate the contribution of each amino acid during the binding of RBD to antibodies [91,92]. The binding free energy for the RBD-Antibody complex was obtained using:

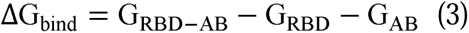

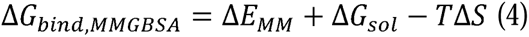

where ΔE_MM_ is total gas phase energy (sum of ΔEinternal, ΔEelectrostatic, and ΔEvdw); ΔGsol is sum of polar (ΔGGB) and non-polar (ΔGSA) contributions to solvation. Here, G _RBD–ANTIBODY_ represent the average over the snapshots of a single trajectory of the complex, G_RBD_ and G_ANTIBODY_ corresponds to the free energy of RBD and antibody respectively.

The polar and non-polar contributions to the solvation free energy is calculated using a Generalized Born solvent model and consideration of the solvent accessible surface area [93]. MM-GBSA is employed to predict the binding free energy and decompose the free energy contributions to the binding free energy of a protein–protein complex on per-residue basis. The binding free energy with MM-GBSA was computed by averaging the results of computations over 1,000 samples from the equilibrium ensembles. First, the computational protocol must be selected between the “single-trajectory” (one trajectory of the complex), or “separate-trajectory” (three separate trajectories of the complex, receptor and ligand). To reduce the noise in the calculations, it is common that each term is evaluated on frames from the trajectory of the bound complex. In this study, we choose the “single-trajectory” protocol, because it is less noisy due to the cancellation of intermolecular energy contributions. Entropy calculations typically dominate the computational cost of the MM-GBSA estimates. Therefore, it may be calculated only for a subset of the snapshots, or this term can be omitted [94,95]. In this study, the entropy contribution was not included in the calculations of binding free energies of the RBD-antibody complexes because the entropic differences in estimates of relative binding affinities are expected to be small owing to small mutational changes and preservation of the conformational dynamics [94,95]. MM-GBSA energies were evaluated with the MMPBSA.py script in the AmberTools21 package [96] and gmx_MMPBSA, a new tool to perform end-state free energy calculations from CHARMM and GROMACS trajectories [97].

### Modeling of Residue Interaction Networks

To analyze protein structures, we employed a graph-based representation where residues are modeled as network nodes, and non-covalent interactions between residue side-chains define the edges [98,99]. This approach captures the spatial and functional relationships between residues, providing insights into the protein structural and dynamic properties. The Residue Interaction Network Generator (RING) program [100–102] was used to generate the initial residue interaction networks from the crystal structures of the protein complexes. We computed the network parameters, such as shortest paths and betweenness centrality, to identify residues critical for communication within the protein structure. The short path betweenness (SPC) of residue *i* is defined to be the sum of the fraction of shortest paths between all pairs of residues that pass through residue *i*:

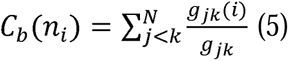

denotes the number of shortest geodesics paths connecting *j* and *k,* and *g _jk_* (*i*) is the number of shortest paths between residues *j* and *k* passing through the node *n_i_.* Residues with high occurrence in the shortest paths connecting all residue pairs have a higher betweenness values. For each node *n,* the betweenness value is normalized by the number of node pairs excluding *n* given as(*N* -1)( *N* - 2) / 2, where *N* is the total number of nodes in the connected component that node *n* belongs to.

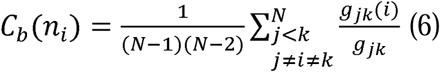

To account for differences in network size, the betweenness centrality of each residue ii was normalized by the number of node pairs excluding ii. The normalized short path betweenness of residue *i* can be expressed as : *g _jk_* is the number of shortest paths between residues *j* and *k*; *g _jk_* (*i*) is the fraction of these shortest paths that pass through residue *i.* Residues with high normalized betweenness centrality values were identified as key mediators of communication within the protein structure network. All parameters were computed using the python package NetworkX [103] and Cytoscape package for network analysis [104–106].

### Network-Based Mutational Profiling of Allosteric Residue Centrality

Through mutation-based perturbations of protein residues we compute dynamic couplings of residues and changes in the short path betweenness centrality (SPC) averaged over all possible modifications in a given position.

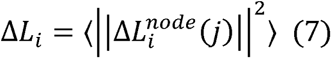

where *i* is a given site, *j* is a mutation and 〈⋯〉 denotes averaging over mutations. 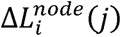 describes the change of SPC parameters upon mutation *j* in a residue node *i*. 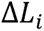 is the average change of ASPL triggered by mutational changes in position *i*. Z-score is then calculated for each node as follows:

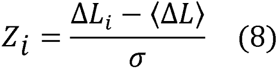

〈Δ*L*〉 is the change of the SPC network parameter under mutational scanning averaged over all protein residues and σ is the corresponding standard deviation. The ensemble-average Z score changes are computed from network analysis of the conformational ensembles of antibody-RBD complexes using 1,000 snapshots of the simulation trajectory.

## Results

### Structural Analysis of the RBD Complexes with Class 4 Antibodies

We first performed structural binding epitope mapping and analysis of closely-related cross-reactive but weakly neutralizing class 4 antibodies CR3022 (group F1, class 4) [31,56,57], EY6A [57] and COVA1-16 [58] (Figure 1, Supporting Information, Tables S1-S3). The CR3022 epitope is typically only accessible when at least two RBDs of the S protein are in the “up” conformation (Figure 1A,B). EY6A and CR3022 bind to the same epitope on the RBD but EY6A binds at about 70 degrees to the perpendicular axis of the α3-helix, which is central to both epitopes (Figure 1C,D) [110]. Both CR3022 and EY6A target residues 368-392 on the RBD, CR3022 interacts with sites 408,427-433, 515-519 while EY6A makes contacts with 411-414, 427-430 (Figures 1,2). The binding epitope for COVA1-16 showed a very similar footprint of residues 368-385, 408, 412-415,427-430 (Figures 1,2).

**Figure 1.**
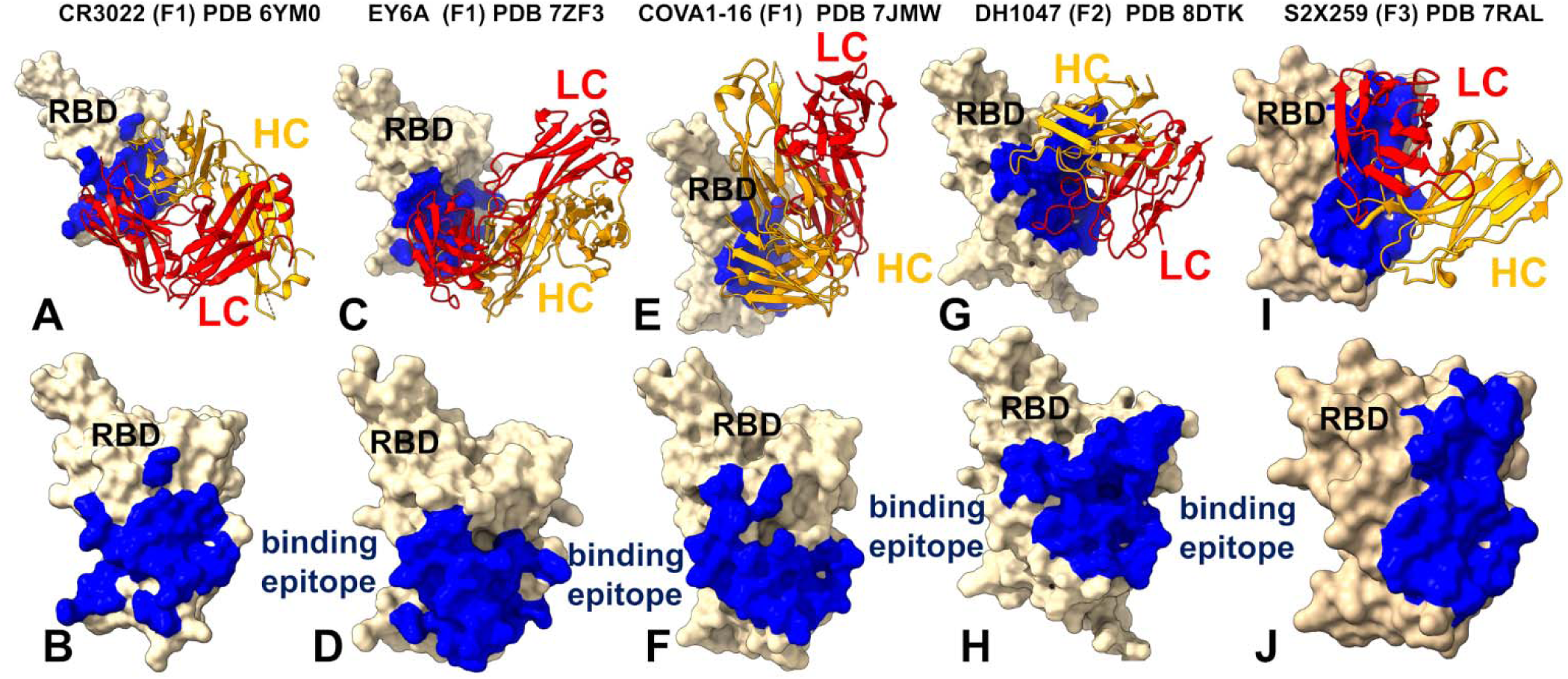
Structural organization of the RBD complexes and binding epitopes for class 4 antibodies. (A) The structure of class 4, group F1 CR3022 with RBD (pdb id 6YM0). The RBD in wheat-colored surface. The heavy chain in orange ribbons, the light chain in red ribbons. (B) The RBD and binding epitope footprint for CR3022. The binding epitope residues are shown in blue surface. (C) The structure of class 4 group F1 antibody EY6A bound with RBD (pdb id 7ZF3). (D) The RBD and binding epitope footprint for EY6A. (E) The structure of class 4 group F1 antibody COVA1-16 bound with RBD (pdb id 7JMW). (F) The RBD and binding epitope footprint for COVA1-16. (G) The structure of class 4 group F2 antibody DH1047 bound with RBD (pdb id 8DTK). (H) The RBD and binding epitope footprint for DH1047. (I) The structure of class 4 group F3 antibody S2X259 bound with RBD (pdb id 7RAL). (D) The RBD and binding epitope footprint for S2X259.

**Figure 2.**
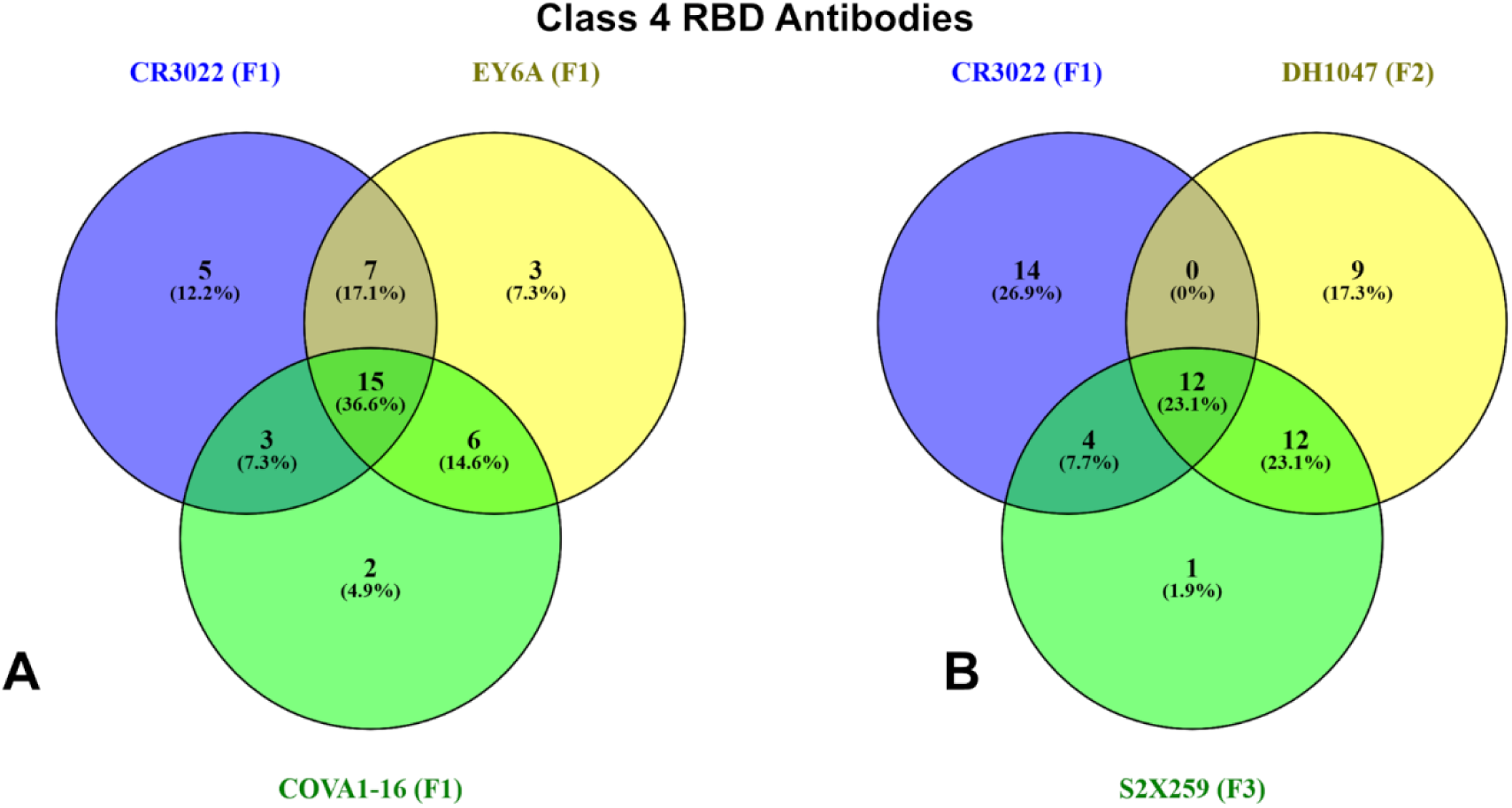
Overlap of Binding Hotspots Among Class 4 RBD Antibodies Across Groups F1 and F2. (A) Venn diagram illustrates the overlap in binding hotspots among group F1 class 4 antibodies CR3022, EY6A, and COVA1-16. The diagram shows the number of residues (and their percentage contributions) that are shared or unique to each antibody within this group. For example, CR3022 (F1) shares 15 residues (36.6%) with both EY6A and COVA1-16, while 5 residues (12.2%) are unique to CR3022. (B) Venn diagram comparing the binding hotspots between group F1 antibody CR3022 (F1) and group F2 antibody DH1047 (F2). This comparison highlights the transition from indirect allostery to partial steric hindrance as observed in group F2. Notably, DH1047 (F2) exhibits a larger set of unique binding hotspots (14 residues, 26.9%) compared to its overlap with CR3022 (4 residues, 7.7%), reflecting its more direct engagement with the ACE2 interface. The green circle represents the overlapping residues common to all three groups, indicating conserved interactions across these antibodies. The percentages indicate the relative contribution of each subset of residues to the total binding hotspots identified for each antibody.

Structural mapping of the binding epitopes for these three antibodies highlighted the same mode of binding and small antibody-specific variations around the cryptic site (Figure 1A-F). It is evident that CR3022 makes contacts with more residues in the binding site, particularly residues 515-519 (Figure 1A,B). However, our analysis strongly suggests that in addition to CR3022, both EY6A and COVA1-16 are class 4 antibodies that can be attributed to the same group F1 (Figure 1C-F).

Members of the F1 group, such as CR3022, EY6A and COVA1-16 require a wide open RBD to engage and do not directly block ACE2, therefore displaying weak neutralizing activities. DH1047 antibody interacts with specific regions of the RBD, including those known to be mobile, through its heavy-chain complementarity-determining region 3 (HCDR3) and light-chain complementarity-determining region 1 (LCDR1) and LCDR3 (Figure 1G,H). By directly engaging with these areas, the antibody can restrict their movement. Structural analysis reveals that the binding of DH1047 to the RBD creates steric hindrance and reaches towards so-called “right shoulder” of the RBD [107], making some partial overlap the ACE2 binding site. The notable extension of the binding epitope for group F2 DH1047 is formation of multiple contacts with R408 and additional interactions with residues 500-508 that overlap with the ACE2 binding site (Figure 1G,H, Supporting Information, Table S4). For group F3 S2X259 we observe even further movement of the epitope to the right shoulder, towards the ACE2 binding site, making contacts with residues 377-385, 404-408, 501-508 (Figure 1I,J, Supporting Information, Table S5).

Group F3 antibodies such as S2X259 reach further towards the ACE2-binding site and interact with appreciable number of ACE2-binding residues therefore directly competing with ACE2 (Figure 1I,J). Interestingly, the key interacting sites D405, R408, V503, G504 and Y508 for group F3 S2X259 (Figure 1I,J, Supporting Information, Table S5) can enable interference with ACE2 but also are associated with the major escape sites for this group of antibodies [26,27,37,38]. Among these, Y501 and H505 are especially important for ACE2 engagement, and by directly interacting with them, S2X259 effectively competes with the host receptor for RBD binding. Because these residues are indispensable for viral entry, they are under strong evolutionary constraint, making them poor candidates for immune escape mutations. S2X259 and other group F3 antibodies are engaged in interactions with peripheral residues D405, and R408 that can mutate and serve as immune escape positions [26,27]. This analysis emphasizes the dual nature of the S2X259 epitope where a central, conserved core which is arguably resistant to mutational drift due to its functional importance, flanked by more flexible peripheral regions that may accommodate escape mutations under selective pressure. Structural analysis reveals that all class 4 antibodies engage a central hydrophobic core of the RBD, accessible only when at least two RBDs adopt the “up” conformation. However, subtle differences in orientation and contact footprint distinguish between each subgroup.

We also illustrated the overlap in the binding epitope residues for CR3022 of group F1, DH1047 of group F2 and S2X259 of group F3 (Figure 2). All three antibodies belong to class 4. The Venn diagram for group F1 antibodies (Figure 2A) reveals that CR3022, EY6A, and COVA1-16 share a significant number of binding hotspots, indicating a common mode of recognition centered around the deeply conserved hydrophobic core of the RBD. The shared residues (∼15%) highlight the conservation of epitope targeting within this group, which explains their broad reactivity across sarbecoviruses. However, each antibody exhibits unique features. CR3022 engages more residues in the binding site, particularly residues 515–519, suggesting a slightly broader footprint compared to EY6A and COVA1-16. EY6A binds at a similar region but with a slightly different orientation, leading to subtle differences in contact geometry and mutational sensitivity. It is notable from structural mapping of the epitopes that group F2 DH1047 has a significant overlap with group F1 antibodies but begins to extend partly towards the ACE2 binding interface (Figure 2B). These diagrams provide insight into the shared and distinct epitope footprints of class 4 antibodies, highlighting how subtle shifts in binding geometry influence their neutralization mechanisms and escape vulnerabilities.

### Conformational Dynamics of the RBD Complexes with Antibodies Using Coarse-Grained and Atomistic Simulations

We performed multiple CG-CABS and atomistic simulations of the RBD-antibody complexes. The root-mean-square fluctuation (RMSF) profiles provide a detailed view of the dynamic behavior of RBD residues upon antibody binding, highlighting both shared features and notable differences among the antibodies. The RMSF analysis of RBD residues for class 4 antibodies provided some insights into the dynamic behavior of the RBD upon antibody binding. These profiles reveal both shared characteristics and particular features for each antibody (Figure 3A). This central β-sheet core (β1: residues 354–358; β2: residues 376–379; β3: residues 394–403; β4: residues 432–437; β5: residues 452–454; β6: residues 492–494; β7: residues 507–516) and α-helices exhibit low RMSF values across all class 4 antibodies, indicating minimal flexibility in these regions. This reflects the conserved structural integrity of the RBD core, which is critical for maintaining its overall stability. Residues such as 350–360, 375–380, and 394–403 show consistently low fluctuations, underscoring their role in stabilizing the RBD structure regardless of the antibody bound (Figure 3A). Conformational dynamics analysis reveals that group F1 antibodies engage a deeply conserved, cryptic epitope located in the core of the RBD, accessible only when at least two RBDs adopt the “up” conformation. These antibodies stabilize the central β-sheet structure of the RBD — composed of residues such as β1–β7 — reducing local flexibility and reinforcing structural integrity. However, their binding also increases mobility in the receptor-binding motif (RBM) loop (residues ∼470–490), promoting greater conformational sampling and facilitating access to the cryptic site. This redistribution of rigidity and flexibility appears to underpin their neutralization mechanism: rather than directly blocking ACE2, they modulate the RBD’s conformational equilibrium, indirectly impairing receptor engagement by favoring non-productive open states. RMSF profiles confirm this pattern, showing a stabilized core with elevated fluctuations in the RBM loop for all F1 antibodies. The increased flexibility in this region may help expose the epitope but also contributes to the relatively weak neutralization potency observed in these antibodies. Importantly, this allosteric modulation aligns with experimental evidence showing altered ACE2 kinetics upon CR3022 binding — slowing association and accelerating dissociation — suggesting that neutralization arises not from physical occlusion, but from long-range dynamic effects [57]. Group F2 antibody DH1047 exhibits a more direct influence on the ACE2 interface compared to F1 antibodies, while still engaging the conserved RBD core. RMSF analysis shows minor increases in flexibility in residues 450–470, coupled with reduced mobility in the RBM-containing loop (∼470–490), indicating a shift in dynamic control toward stabilizing the ACE2-contacting regions (Figure 3B). This stabilization likely supports a hybrid mechanism of action, where DH1047 exerts partial steric hindrance on ACE2 binding while maintaining allosteric effects via interactions with structurally conserved sites. Cryo-EM and simulation data suggest that DH1047 binding restricts the RBD’s ability to transition between conformations, locking it into states less favorable for viral entry. This dual mode — combining dynamic modulation and localized interference — enhances neutralization potency relative to group F1, though at the cost of increased sensitivity to mutations. Notably, Omicron BA.2 mutations such as T376A and R408S severely compromise DH1047 binding, highlighting how increased proximity to the receptor interface correlates with escape vulnerability [26,27]. Overall, DH1047 may represent an intermediate evolutionary and dynamic adaptation among class 4 antibodies — leveraging both allostery and direct interference to achieve stronger neutralization without fully overlapping with the ACE2 footprint.

**Figure 3.**
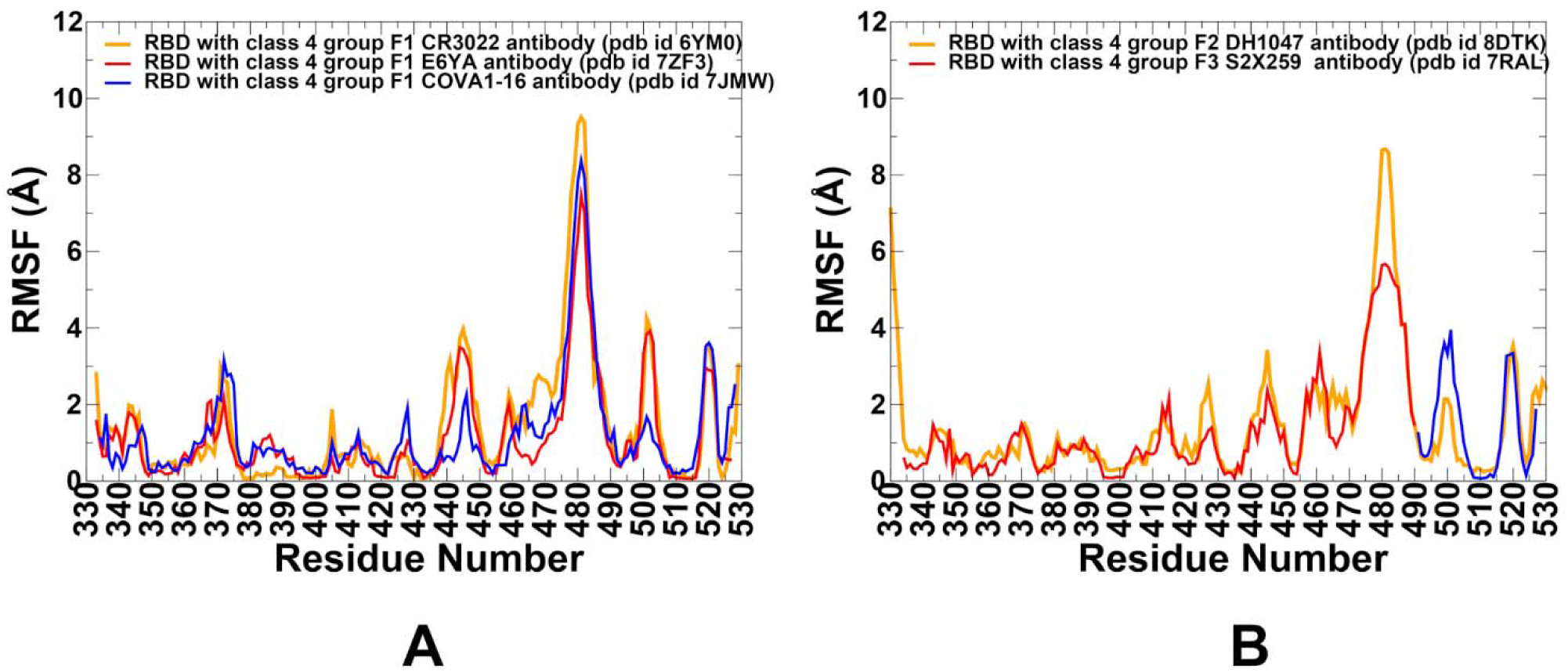
RMSF Profiles of RBD Residues Upon Binding with Class 4 Antibodies. (A) RMSF profiles for residues of the RBD in complexes with group F1 class 4 antibodies. The figure compares the dynamic behavior of the RBD upon binding to CR3022 (PDB ID: 6YM0, orange), EY6A (PDB ID: 7ZF3, red), and COVA1-16 (PDB ID: 7JMW, blue). Residues within the central β-sheet structure (e.g., residues 354–358, 376–379, 394–403, 432–437, 452–454, 492–494, and 507–516) exhibit low RMSF values across all three antibodies, indicating minimal flexibility. Residues ∼470–490 show elevated RMSF values, particularly for group F1 antibodies, reflecting enhanced local flexibility in this region. (B) RMSF profiles for residues of the RBD in complexes with group F2 (DH1047) and group F3 (S2X259) class 4 antibodies. Group F2 (DH1047, PDB ID: 8DTK, orange) : Residues 450–470 exhibit moderately increased mobility, while the 470–490 loop shows reduced flexibility compared to group F1 antibodies. Group F3 (S2X259, PDB ID: 7RAL, blue) : Similar to DH1047, S2X259 also displays reduced flexibility in the 470–490 loop, consistent with its more direct engagement of the ACE2 interface.

The group F3 antibody S2X259 demonstrates the most pronounced impact on RBD dynamics, reflecting its direct competition with ACE2. RMSF profiles show significantly reduced flexibility in the RBM loop (∼470–490), consistent with strong binding that locks the RBD into a conformation incompatible with efficient ACE2 engagement (Figure 3B). These shifts suggest that both DH1047 and S2X259 modulate RBD dynamics in distinct ways, subtly reshaping the conformational landscape to enhance direct competition with ACE2. The observed stabilization of key interface residues likely contributes to their increased neutralization potency compared to earlier group F1 antibodies, offering a mechanistic link between dynamic modulation and receptor blocking. To sum up, the RMSF profiles reveal a progressive shift in conformational dynamics across class 4 antibodies from groups F1 to F3. Group F1 antibodies stabilize the core while increasing flexibility in the RBM loop, promoting allosteric interference with ACE2 binding (Figure3, Table 1). In contrast, groups F2 and F3 exhibit reduced flexibility in the RBM loop, reflecting their closer proximity to the ACE2 interface and stronger interactions with critical residues involved in receptor binding (Table 1). These results may underscore the mechanistic evolution of neutralization strategies among class 4 antibodies, transitioning from indirect allostery (F1) to partial steric hindrance (F2) and direct competition with ACE2 (F3).

**Table 1.** Summary of the Structural and Dynamics Analysis for Class 4 Antibodies.

|  | Group F1 antibodies | Group F2 antibodies | Group F3 antibodies |
| --- | --- | --- | --- |
| **Binding Site** | Core RBD, cryptic | Partial overlap + | Further shift toward |
|  |  | ACE2 interface | ACE2 interface |
| **ACE2 Competition** | No | Partial | Yes |
| **Key Interactions** | Leu445, Phe486,<br>Tyr505 | R408, 500–508 | D405, R408, V503,<br>G504, Y508 |
| **Escape Mutations** | 383–386, 390, 391 | 408, 500–508 | 501, 505 |
| **RMSF Profile** | Stabilized core, flexible<br>470–490 loop | Moderate flexibility<br>in 450–470 | Reduced flexibility in<br>470–490 |
| **Neutralization**<br>**Mechanism** | Allosteric | Partial steric<br>hindrance | Direct competition<br>with ACE2 |

### Mutational Profiling of Antibody-RBD Binding Interactions Interfaces Reveals Molecular Determinants of Immune Sensitivity and Emergence of Convergent Escape Hotspots

Using the conformational ensembles of the RBD-antibody complexes, we embarked on structure-based mutational analysis of the S protein binding with antibodies. To provide a systematic comparison, we constructed mutational heatmaps for the RBD interface residues of the S complexes with class I antibodies. Mutational scanning studies of RBD–antibody complexes provides insight into the epitope sensitivity and escape vulnerability of class 4 group F1 antibodies — CR3022, EY6A, and COVA1-16. These analyses, derived from conformational ensembles and DMS datasets, reveal a consistent pattern: while these antibodies target a highly conserved core epitope, their neutralizing activity is constrained by specific mutation-induced disruptions. Mutational profiling of CR3022 binding identifies a set of key interface residues that are essential for antibody recognition. F377, C379, Y380, T385 emerged as dominant hotspots (Figure 4A). Additional contributions come from K378, G381, V382, S383, P384, L390, and F392, which also play supportive roles in maintaining epitope integrity. Our results are consistent with the DMS data revealing that residues 383–386, 390, and 392 represent major escape hotspots for CR3022 and other F1-class antibodies. These residues lie at the edge of the epitope, suggesting that immune pressure drives mutations here to evade antibody recognition without compromising viral function. EY6A binds to a nearly identical region on the RBD as CR3022 (Figures 1,2) yet exhibits somewhat distinct mutational sensitivity profile reflecting differences in orientation and contact geometry. The heavy chain of CR3022 features hotpots in positions I30, T31, Y32, W33 (Figure 4B). Mutational profiling revealed that EY6A engages similar dominant residues — including F377, C379, Y380, T385, and S383 — confirming that both antibodies recognize a shared structural motif. However, EY6A also makes weaker interactions with P412 and G413, which are located slightly further from the central hotspot (Figure 4C). This broader interaction profile may offer modest resilience to certain escape mutations, although it remains sensitive to changes at key peripheral sites. The heavy chain of EY6A shows strong hotspot contributions from Y59, W104, V105, and Y106, underscoring the importance of aromatic stacking and van der Waals forces in stabilizing this complex (Figure 4D).

**Figure 4.**
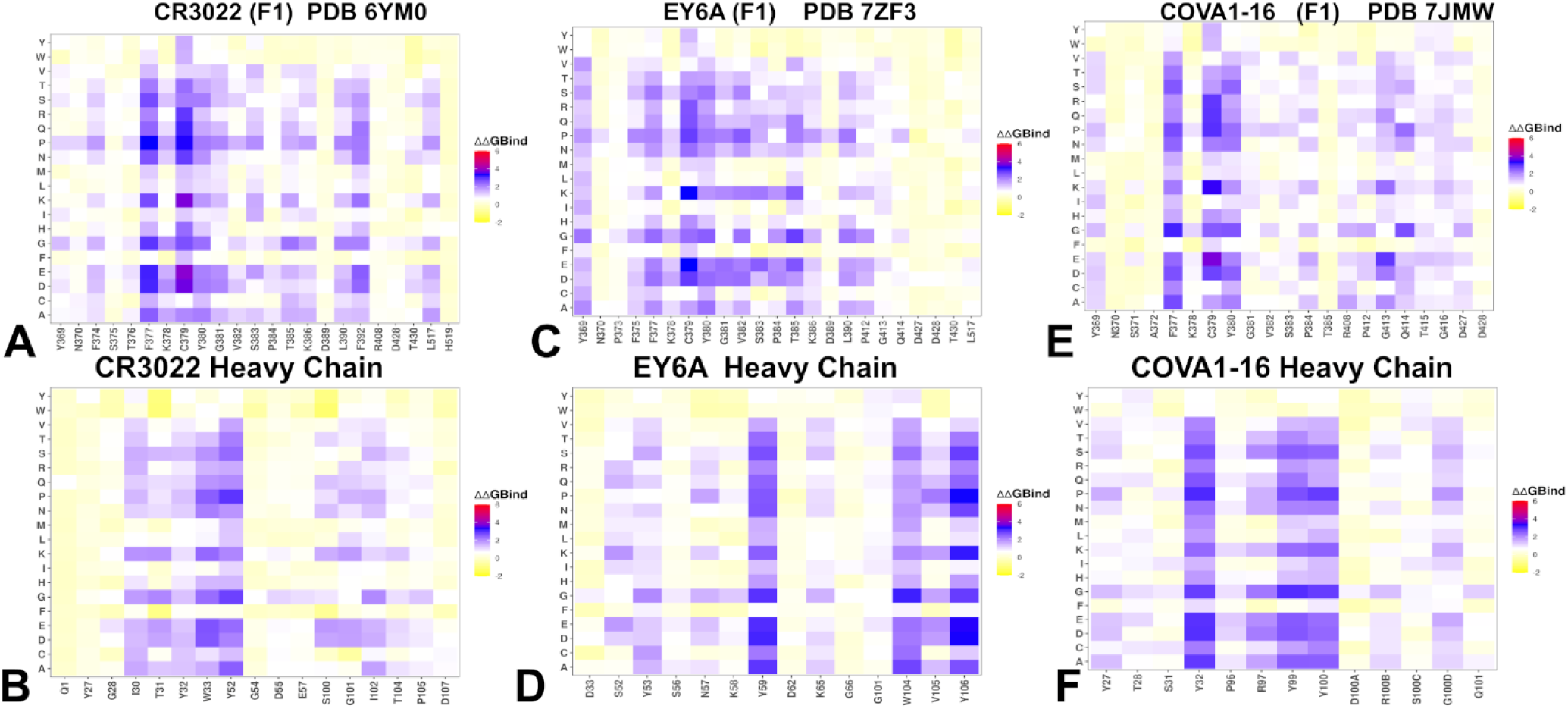
Ensemble-based dynamic mutational profiling of the RBD intermolecular interfaces in the RBD complexes. Mutational heatmaps for RBD complex with group F1 CR3022 antibody (A,B), the RBD complex with group F1 E6YA antibody (C,D) and the RBD complex with group F1 COVA1-16 antibody (E,F). The mutational scanning heatmaps are shown for the interfacial RBD residues and interfacial heavy chain residues of respective class 4 group F1 antibodies. The heatmaps show the computed binding free energy changes for 20 single mutations of the interfacial positions. The standard errors of the mean for binding free energy changes using randomly selected 1,000 conformational samples (0.12-0.18 kcal/mol) obtained from the atomistic trajectories.

For COVA1-16, mutational analysis confirms a binding mode overlapping with CR3022 and EY6A, targeting residues 368–385, 408, 412–415, and 427–430, with major contributions from F377, K378, C379, and Y380 (Figure 4E). Minor hotspots such as R408 and G413 were also identified, suggesting that COVA1-16 extends its influence closer to the ACE2 interface than CR3022 or EY6A. Despite this, it retains the hallmark features of group F1 antibodies — broad reactivity due to targeting a structurally conserved RBD core, and vulnerability to mutations affecting epitope integrity or solvent exposure. Its heavy chain interactions are dominated by Y32, Y99, and Y100, highlighting a unique residue-level specificity that may shape its conformational adaptability and dynamic coupling with the RBD (Figure 4F). Taken together, these mutational scanning results highlight a trade-off between conservation and conformational dependency in group F1 antibodies. While their targeting of a deeply conserved RBD core ensures broad sarbecovirus reactivity, their reliance on rare “up” conformations and sensitivity to edge mutations limits their neutralization potency, especially against evolving variants.

The mutational scanning analysis of group F2 antibody DH1047 and group F3 antibody S2X259 provides key insights into how these class 4 antibodies engage the RBD and respond to immune-evading mutations. While both target epitopes that overlap with the ACE2 interface, they do so with distinct energetic and functional consequences. For DH1047, mutational profiling identifies a set of critical RBD residues — including Y369, F374, T376, F377, C379, Y380, R408, V503, Y505, and Y508 — that are essential for antibody recognition. Among these, T376, R408, V503, and Y508 emerge as major destabilization sites, where single-point mutations such as T376A (ΔΔG = 2.15 kcal/mol), R408A/S (ΔΔG = 2.35–2.44 kcal/mol), and Y508H/N significantly impair binding (Figure 6A).

**Figure 5.**
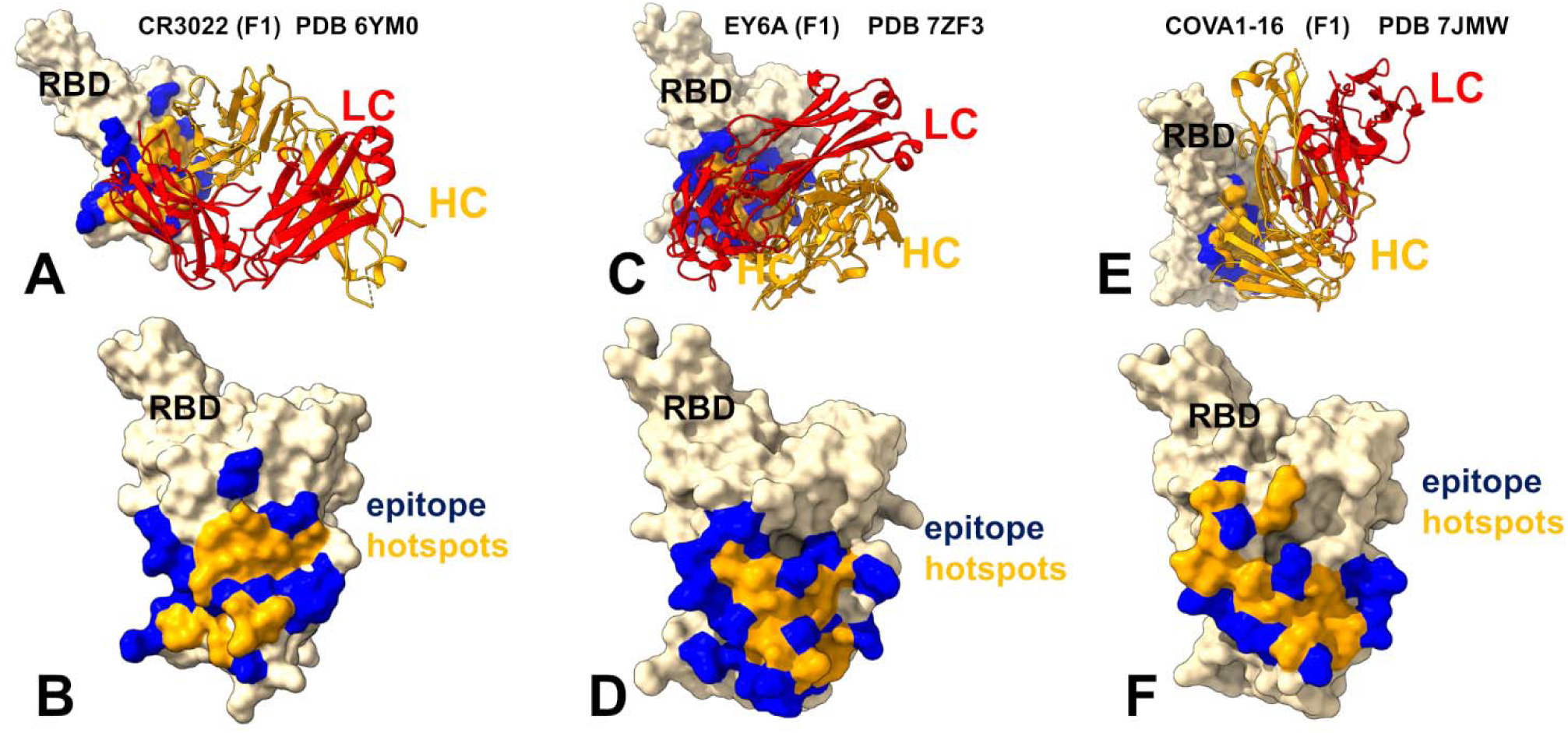
Structural and Epitope Mapping of Group F1 Class 4 Antibodies Binding to the RBD. The structures and epitope hotspots for three representative group F1 class 4 antibodies — CR3022 (PDB ID: 6YM0), EY6A (PDB ID: 7ZF3), and COVA1-16 (PDB ID: 7JMW). The three-dimensional structures of the RBD–CR3022 complex (A), RBD-EY6A complex (C) and COVA1-16/RBD (F). The RBD is depicted in wheat-colored surface, with the heavy chain (HC) of the antibody in orange and the light chain (LC) in red. The binding epitope residues are shown in blue surface and the RBD binding hotspots are shown in orange surface. Panels B, D, and F provide a detailed view of the RBD, the epitope and binding hotspots for CR3022 (PDB ID: 6YM0), EY6A (PDB ID: 7ZF3), and COVA1-16 (PDB ID: 7JMW) respectively. The epitope sites are highlighted in blue surface, and binding hotspots are in orange surface. These panels reveal that while all three antibodies target a deeply conserved hydrophobic core within the RBD, there are subtle differences in their contact footprints and residue-specific interactions, reflecting minor variations in orientation and binding geometry.

**Figure 6.**
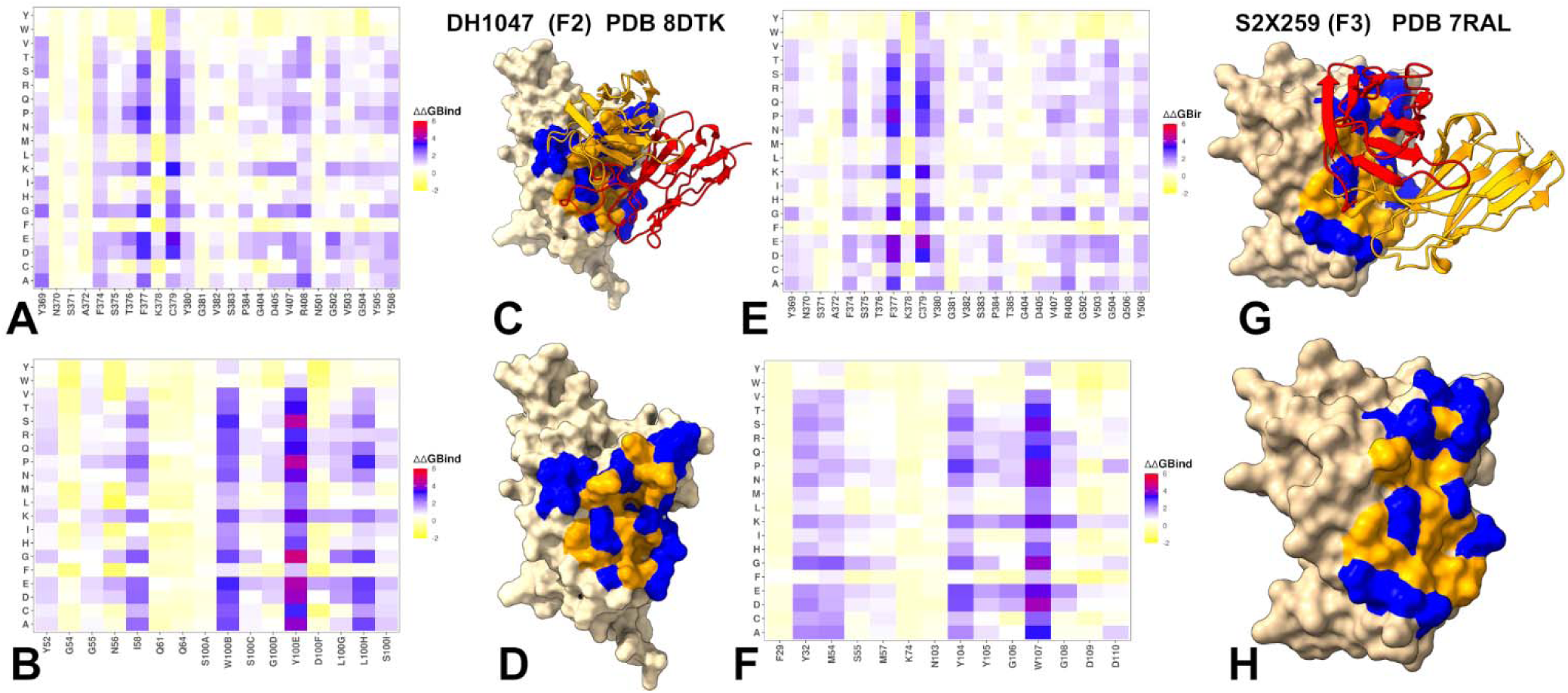
Mutational Heatmaps and Epitope Mapping of Group F2 and F3 of Class 4 Antibodies Binding to the RBD. Mutational heatmaps of the RBD binding interface residues for RBD complex with group F2 DH1047 antibody (A) and mutational heatmap of the heavy chain of DH1047 (B). The three-dimensional structures of the DH1047 complex with RBD (C), and a detailed view of the RBD, the epitope and binding hotspots for DH1047 (D) (PDB ID: 8DTK). Mutational heatmaps of the RBD binding interface residues for RBD complex with group F3 S2X259 antibody (E) and mutational heatmap of the heavy chain of S2X259 (F). The three-dimensional structures of the S2X259 complex with RBD (C), and a detailed view of the RBD, the epitope and binding hotspots for S2X259 (D) (PDB ID: 7RAL). The RBD is in wheat-colored surface. The epitope sites are highlighted in blue surface, and binding hotspots are in orange surface.

The emergence of these mutations highlights a growing escape risk associated with proximity to the ACE2 interface, despite the functional constraints on many of these residues. These findings indicate that even though the core RBD region remains conserved, certain peripheral residues under immune pressure are mutable, especially those near the ACE2-binding motif. Importantly, the heavy chain residue Tyr100D in DH1047 forms extensive hydrogen bonds with Ser371, Thr376, Phe377, and Tyr369, reinforcing the idea that this region plays a critical role in stabilizing the antibody–RBD complex (Figure 6B). However, the presence of Omicron BA.1/BA.2 mutations (S371L, S373P, S375F) indirectly compromises binding by altering the local conformation or solvent exposure of nearby hotspot residues, further highlighting the network-level sensitivity of DH1047 to changes in the RBD landscape. Structural mapping of the DH1047 complex with RBD and the footprint of the binding hotspots on the RBD (Figure 6C,D) illustrates a broadly distributed allocation of the binding hotspots. Notably the partial overlap with the ACE2 binding site does not involve strong binding hotspots suggesting that these interactions can modulate the binding and immune escape patterns for this group of antibodies.

Group F3 antibody S2X259 engages the RBD through an even more direct overlap with the ACE2 footprint, interacting strongly with D405, R408, V503, G504, and Y508 — all of which are key contact points for host receptor engagement (Figure 6E,F).. This proximity enables direct competition with ACE2, resulting in stronger neutralization potency, but also increases susceptibility to immune escape. Mutational data show that substitutions such as D405N, R408S, and T376A severely disrupt binding, consistent with their high energetic contributions to the interaction (Figure 6E). R408 and T376 remain mutable under selective pressure, making them high-risk escape positions. The emergence of T376A and R408S in Omicron BA.2 exemplifies how immune pressure drives mutations at the periphery of functionally constrained regions, compromising antibody binding while preserving viral fitness [26,27,37,38]. This analysis underscores the trade-off between neutralization strength and escape resistance, driven by the progressive shift from allostery toward steric interference across class 4 antibodies.

### MM-GBSA Analysis of the Binding Energetics for Class 4 Antibodies Complexes

The MM-GBSA analysis offers a detailed energetic dissection of how class 4 antibodies engage the RBD revealing distinct patterns of interaction that reflect their neutralization mechanisms, escape vulnerabilities, and evolutionary positioning within the antibody classification system. By decomposing the total binding free energy into its van der Waals (VDW) and electrostatic (ELE) components, we gain insight into the molecular determinants of binding affinity, and how mutations at key positions can disrupt these interactions, leading to immune evasion. For group F1 antibodies CR3022, EY6A, and COVA1-16, the MM-GBSA results highlight a strong reliance on hydrophobic interactions, particularly involving residues 377-382, which form a tightly packed core critical for stabilizing the antibody–RBD complex. These residues are part of the central β-sheet structure of the RBD and are highly conserved across sarbecoviruses, explaining the broad reactivity of this group (Figure 7A,B). In contrast, electrostatic contributions are modest and distributed, with K378 and D428 showing favorable interactions that help fine-tune the binding interface but do not dominate energetically (Figure 7C).

**Figure 7.**
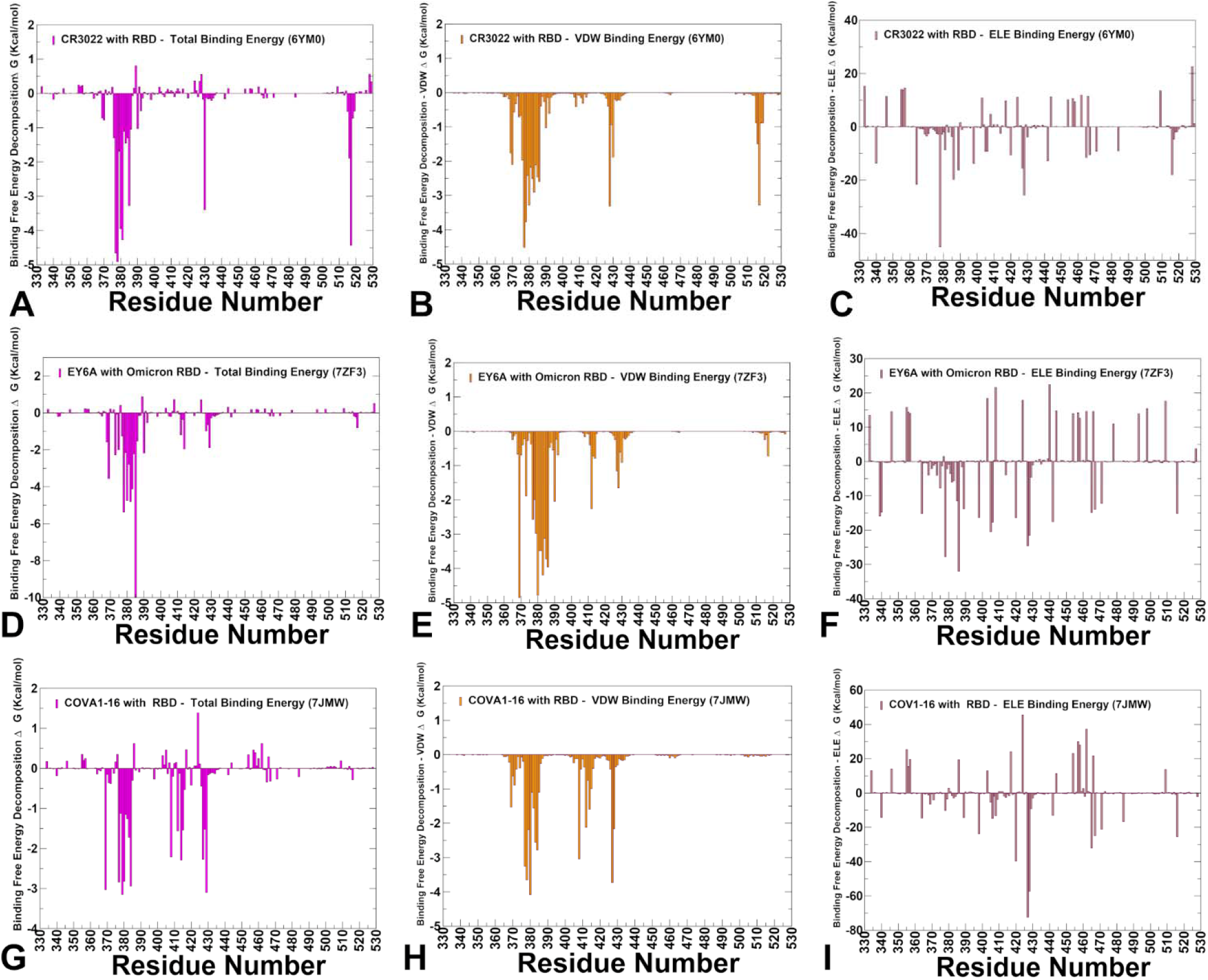
Binding Free Energy Decomposition of Group F1 Antibodies with the RBD Using MM-GBSA Analysis. MM-GBSA binding free energy decomposition for three representative group F1 class 4 antibodies — CR3022, EY6A, and COVA1-16 in complex with RBD. (A) The total binding free energy for CR3022 binding to the RBD, highlighting key residues contributing to overall stability. (B) Van der Waals (VDW) component of the binding free energy, showing dominant contributions from hydrophobic interactions. (C) Electrostatic (ELE) component of the binding free energy, revealing residues critical for charge-based stabilization. ( D) The total binding free energy for EY6A binding to the RBD, emphasizing energetic contributions across the interface. (E) VDW contribution to binding, illustrating the role of nonpolar contacts in stabilizing the complex. (F) ELE contribution to binding, identifying residues involved in electrostatic interactions. (G) The total binding free energy for COVA1-16 binding to the RBD. (H) VDW component of the binding free energy, highlighting residues that mediate hydrophobic interactions. (I) ELE component of the binding free energy, pinpointing residues crucial for electrostatic complementarity. Residue-based binding free energy values are shown for total energies as magenta-colored filled bars, van der Waals contributions as orange-colored bars and electrostatic contributions as light-brown colored bars. MM-GBSA contributions are evaluated using 1,000 samples from MD simulations.

EY6A binds to a very similar region of the RBD as CR3022, yet its energetic profile reveals some specific features. The central core residues 378-385 are the main contributors to binding, with favorable van der Waals interactions playing a dominant role (Figure 7D,E). EY6A engages a slightly broader region than CR3022, including residues P412 and G413, which may offer some resilience against localized mutations (Figure 7D,E). While K378, K386, D405, D427, and D428 exhibit favorable electrostatic contacts (Figure 7F), these interactions are largely offset by unfavorable solvation penalties, resulting in only marginal net stabilization of the complex. This suggests that EY6A relies more heavily on hydrophobic interactions than on electrostatic complementarity, making it potentially more robust against certain polar substitutions, but vulnerable to mutations that alter hydrophobic packing including those seen in Omicron variants.

COVA1-16 presents an energy profile, marked by a more distributed pattern of hotspot residues compared to CR3022 and EY6A. Key contributors include C379, Y369, F377, Y380, P384, along with Y380, D427, and R408, emphasizing the importance of aromatic stacking and aliphatic interactions in stabilizing the complex (Figure 7G,H). Electrostatically, favorable contributions arise from D420, D427, and D428, reinforcing the idea that charged residues play a supportive but secondary role in binding (Figure 7I). This broader footprint may confer some resilience to localized mutations, although it also increases the number of potential escape pathways, particularly at positions 412–415 and 427–428, which are under growing immune pressure. Importantly, the interactions of COVA1-16 extend slightly toward residues partly overlapping with the ACE2 interface, suggesting that its neutralization mechanism may involve a mild competitive component, despite being classified as a class 4 antibody.

One of the most striking insights from this analysis is the delicate interplay between different energetic components that define the binding landscape of group F1 antibodies. In CR3022 and EY6A, hydrophobic interactions dominate, contributing to high conservation and broad reactivity, but also rendering them vulnerable to mutations that destabilize local structure. In contrast, a somewhat wider hotspot distribution and greater involvement of electrostatics found for COVA1-16 antibody suggest a more adaptable binding mode, though one that may be more sensitive to polar or charged mutations. These energetic trade-offs have clear implications for immune evasion. Mutations T376A and R408S, found in Omicron BA.2 can disrupt critical hydrophobic or electrostatic contacts, leading to complete loss of neutralization by these antibodies (Figure 7) Even conserved residues F377 and C379, though rarely mutated due to structural constraints, can be compromised by second-order effects such as loop destabilization or alterations in solvent exposure. From a functional perspective, the MM-GBSA results reinforce that group F1 antibodies rely on a delicate balance of forces — dominated by hydrophobic interactions, supported by electrostatics, and constrained by structural rigidity.

The strongest binding hotspots for group F2 DH1047 include Y505, R408, V503, T375, F377, K378, Y380, C379, T415, and F374 (Figure 8A). These residues form a structurally conserved yet partially exposed interface, highlighting the delicate balance between conservation and accessibility. For this group F2 antibody, residues V503, Y505, T376, S375, Y380, F377, and R408 dominate the hydrophobic contacts, providing the primary stabilizing force for the complex (Figure 8B). While K378, R408, and K386 contribute favorable electrostatics, these interactions are offset by solvation penalties, resulting in only marginal net stabilization (Figure 8C). DH1047 engages critical residues R408, V503, and Y505, which are also essential for ACE2 binding. Notably, DH1047 exhibits a hybrid interaction pattern, combining elements of both group F1 conservation and group F3 interference with ACE2 binding. This suggests that group F2 antibodies may occupy an evolutionary intermediate stage, where increased potency comes at the cost of greater sensitivity to immune pressure, especially in evolving variants.

**Figure 8.**
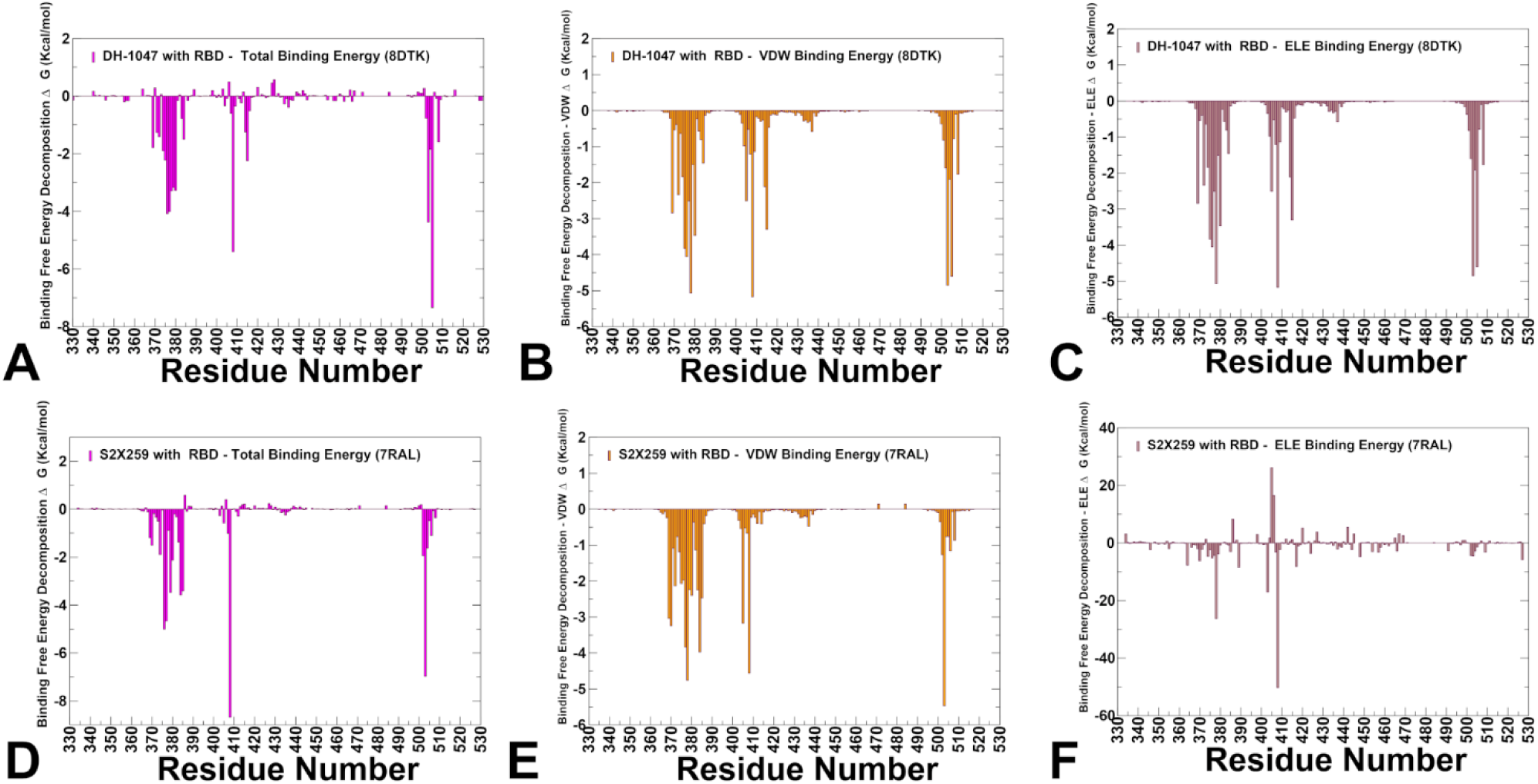
Binding Free Energy Decomposition of Group F2 and F3 Antibodies with the RBD Using MM-GBSA Analysis. MM-GBSA binding free energy decomposition for group F2 DH1047 and group F3 S2X259 in complex with RBD. (A) The total binding free energy for DH1047 binding to the RBD, highlighting key residues contributing to overall stability. (B) Van der Waals (VDW) component of the binding free energy, showing dominant contributions from hydrophobic interactions. (C) Electrostatic (ELE) component of the binding free energy, revealing residues critical for charge-based stabilization. ( D) The total binding free energy for S2X259 binding to the RBD (E) VDW contribution to binding, illustrating the role of nonpolar contacts in stabilizing the complex. (F) ELE contribution to binding, identifying residues involved in electrostatic interactions. Residue-based binding free energy values are shown for total energies as magenta-colored filled bars, van der Waals contributions as orange-colored bars and electrostatic contributions as light-brown colored bars. MM-GBSA contributions are evaluated using 1,000 samples from MD simulations.

Hence, MM-GBSA profile of DH1047, a representative group F2 class 4 antibody, reveals a transition in energetic strategy, with increased involvement of ACE2-proximal residues such as R408, V503, and T376. These sites contribute strongly through hydrophobic contacts, while K378 and R408 also provide favorable electrostatic contributions (Figure 8C). S2X259, a representative group F3 antibody, exhibits a binding mode that extends even closer to the ACE2 interface, which may effectively transition mechanistically from allosteric effects on ACE2 binding toward direct competitive inhibition. The MM-GBSA results highlighted several key features. We found that residues V503, R408, P384, F377, and D405 form robust hydrophobic interactions, anchoring the antibody firmly to the RBD (Figure 8D,E). The favorable electrostatic interactions are observed at R408, K378, R403, D389, and K417, further stabilizing the complex (Figure 8F). The combination of strong hydrophobic and electrostatic interactions creates a robust binding interface that overlaps significantly with the ACE2 footprint, particularly at R408, V503, and G502, which are among the most critical residues for viral entry. The MM-GBSA results suggest that neutralization potency of S2X259 stems from its ability to engage both hydrophobic cores and charged interfaces, thereby combining structural conservation with functional interference. However, this proximity to ACE2 also introduces escape vulnerabilities, particularly at D405 and R408, where substitutions such as D405N and R408S have been shown to significantly reduce binding. Mutational scanning supports these findings, indicating that while only a few residues (e.g., D405, R408, V503/G504) represent major points of vulnerability, they are biochemically significant — often requiring dramatic amino acid changes to evade antibody recognition. Nonetheless, the emergence of T376A, D405N, and R408S mutations in Omicron BA.2 highlights how selective pressure can still drive escape even at functionally constrained sites [26,27]. MM-GBSA calculations reveal that these interactions act synergistically, creating a robust binding interface that enables direct steric competition with ACE2. This explains the highest neutralization potency observed among class 4 antibodies. However, it also introduces significant escape risks, particularly at R408 and D405, where substitutions severely reduce binding affinity. These findings are fully consistent with DMS data which identified R408S and D405N as high-impact escape mutations. While some of these residues are evolutionarily constrained due to their essential role in viral entry, others remain mutable under selective pressure, reflecting the functional duality of this region: structural conservation vs. immunogenic exposure.

When comparing the energetic profiles and escape vulnerabilities across group F1 (CR3022, EY6A, COVA1-16), group F2 (DH1047), and group F3 (S2X259), an interesting evolutionary trajectory emerges reflecting a progression from deeply cryptic, allosteric engagement (F1), through partial overlap with ACE2 and enhanced hydrophobic/electrostatic synergy (F2), to direct receptor competition with strong energetic coupling (F3. From this comparative view, we observe a gradual shift in energetic strategy. Group F1 antibodies rely on deeply conserved hydrophobic cores, offering broad reactivity but low potency due to low accessibility and indirect neutralization mechanisms. Group F2 antibodies begin to leverage electrostatic complementarity, enhancing affinity and partial receptor blocking, but at the cost of increased escape susceptibility. Group F3 antibodies achieve the strongest binding synergy, combining hydrophobic stability with electrostatic reinforcement, allowing them to effectively compete with ACE2 — albeit with greater vulnerability to mutations at key receptor-contacting residues. Our results enable energetic rationalization why group F1 class 4 antibodies excel in breadth and conservation but suffer from limited access and weak binding. In contrast, group F2 antibodies offer a balanced middle ground, with improved potency and moderate escape risk. Finally, group F3 antibodies deliver strong neutralization, but face evolving escape challenges, particularly at R408, D405, and T376.

### Exploring Allosteric Binding Pathways Using Dynamic Network Analysis

We used the ensemble-based network centrality analysis and the network-based mutational profiling of allosteric residue propensities that are computed using the topological network parameters particularly the SPC and Z-Score of the ASPL in the network where we compute changes in the metrics averaged over all possible modifications in a given position for all RBD residues in the complex. Through ensemble-based averaging over mutation-induced changes in the network parameters we identify positions in which mutations on average cause network changes. Allosteric hotspots are identified as residues in which mutations incur significant perturbations of the global residue interaction network that disrupt the network connectivity and cause a significant impairment of network communications and compromise signaling.

The network analysis of allostery for group F1 antibodies CR3022, EY6A, and COVA1-16 reveals a unified yet subtly differentiated mechanism by which these antibodies engage the RBD. By leveraging dynamic simulations, mutational scanning, and energetic decomposition, we have uncovered key insights into how these antibodies stabilize the RBD core while inducing subtle shifts in conformational flexibility, particularly within the RBM loop (∼470–490). These findings underscore the modular nature of RBD allostery, where conserved structural motifs form the backbone of antibody recognition, while peripheral residues mediate functional consequences such as indirect interference with ACE2 binding. Across all three group F1 antibodies, the central hydrophobic core of the RBD remains a conserved interaction hub, stabilized by residues forming β-strands (β1–β7) and α-helices. This core exhibits minimal fluctuations, reflecting its critical role in maintaining overall RBD integrity. However, the 470– 490 loop, which contains the RBM, shows elevated RMSF values, indicating increased local flexibility upon antibody binding. This redistribution of rigidity and mobility suggests that group F1 antibodies induce allosteric effects by stabilizing the central RBD core while promoting open conformations favorable for epitope exposure (Figure 9).

**Figure 9.**
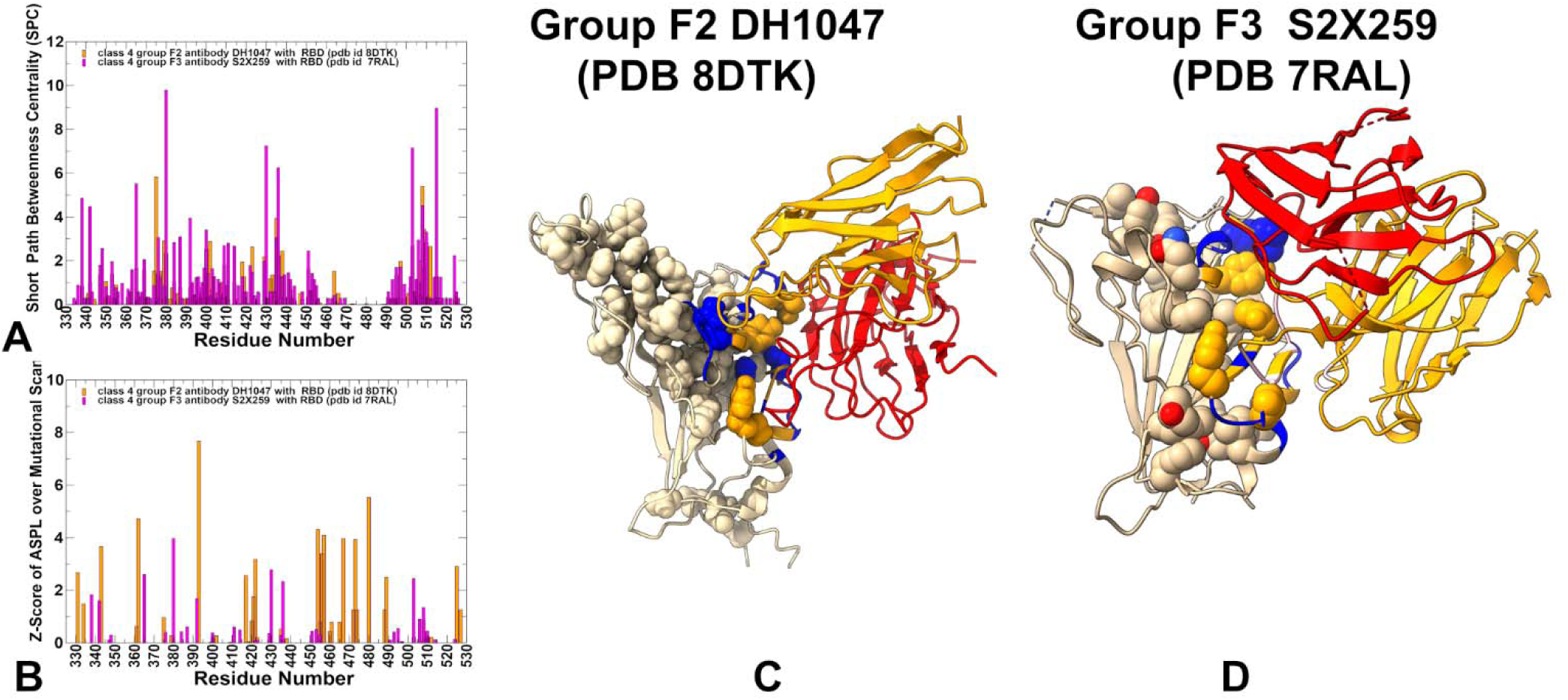
The ensemble-averaged SPC centrality (A) and the average Z-score of ASPL over mutational scan (B) for the RBD residues for the class 4 group F1 antibody complexes : CR3022 with RBD, pdb id 6YM0 (in orange filled bars), EY6A with RBD, pdb id 7ZF3 (in magenta filled bars) and COVA1-16 with RBD, pdb id 7JMW ( in green filled bars). (C) Structural mapping of allosteric network centers for class 4 group F1 CR3022 antibody with RBD. (D) Structural mapping of allosteric network sites for class 4 group F1 COVA1-16 antibody with RBD. The heavy chain in orange ribbons, the light chain in red ribbons. The binding epitope residues are shown in blue surface, and binding hotspots are in orange and allosteric residue interaction network with high SPC and Z-score ASPL values in wheat-colored spheres.

We first analyzed the distribution of the SPC and Z-score ASPL parameter for class 4 group F1 CR3022, EY6A and COVA1-16 antibodies (Figure 9A,B). The high centrality RBD sites for CR3022 include core residues 431,380,396,377,423,436,374,381,355,392,398. Among sites with appreciable Z-Score ASPL values are also RBD positions involved in ACE2 binding such as RBD sites 451,495,497,439,443,506 (Figure 9A,B). A number of mediating network positions for EY6A included RBD core residues 365,338, 342,369, 400,410,407 along with residues closer the ACE2 binding site (495,453, 493,506). For COVA1-16 antibody, the high centrality positions also include RBD core residues 414, 418, 355,436, 396,341,338,379, 393, 397 as well as some residues located near ACE2 binding site ( residues 493, 406, 495). Across all three antibodies, the central RBD core emerges as a highly connected hub in the residue interaction network. For CR3022, high-centrality residues include 431, 380, 396, 377, 423, 436, 374, 381, 355, 392, and 398 — many of which are deeply conserved and structurally critical. These positions form a rigid scaffold that underpins the overall stability of the RBD and serves as a conduit for allosteric signals originating from the cryptic epitope. Similarly, EY6A and COVA1-16 also highlight core residues as key mediators of network connectivity. EY6A engages residues such as 365, 338, 342, 369, 400, 410, and 407, while COVA1-16 identifies 414, 418, 355, 436, 396, 341, 338, 379, 393, and 397 as high-centrality nodes (Figure 9A,B). Notably, several of these residues overlap with those highlighted by CR3022, reinforcing the idea that a shared structural framework exists across group F1 antibodies, even if each antibody approaches it with slight variations in orientation or contact profile. Importantly, all three antibodies appear to influence residues near the ACE2 interface, including 451, 495, 497, 439, 443, and 506, through indirect allosteric coupling rather than direct steric interference. This suggests that despite not directly overlapping with the ACE2-binding motif, group F1 antibodies can still exert functional consequences on receptor engagement, mediated via long-range dynamic reorganization of the RBD. Structural mapping of the high centrality sites in the CR3022 complex (Figure 9C) and COVA1-16 complex (Figure 9D) displayed a dense interaction network that produced rigidification of the RBD central core engaging “the right shoulder” of the RBD [107].

An intriguing and consistent finding across all three antibodies is the lack of high-centrality nodes in the “left shoulder” of the RBD ( residues 450-494) [107] including flexible RBM region. This segment, known for its conformational variability, appears to be only weakly integrated into the core allosteric network, suggesting that its dynamic behavior is largely independent of the structured core. In other words, the allosteric RBD network is weakly coupled to stochastic movements of 450-495 region that reflects the increased mobility of this region. These findings are consistent with the dynamic analysis showing that group F1 antibodies can induce stabilization of the central RBD core coupled with the increased flexibility in the 470–490 loop, thus leading to partial redistribution of the rigidity-flexibility patterns. The increased variations in the RBM regions can ensure highly open RBD conformation in which the cryptic site is available for binding thus enabling allosteric interference with the efficient ACE2 binding. In essence, the allosteric signal propagates through the rigid core, subtly reshaping the dynamical landscape of the entire RBD, and thereby influencing viral entry potential from a distance.

One of the most striking findings from this comparative analysis is the common effect these antibodies have on RBD flexibility. The data consistently show that antibody binding stabilizes the rigid RBD core while increasing mobility in the RBM loop region. The increased conformational diversity in this region may disrupt the precise alignment required for efficient ACE2 binding, subtly altering the energetic landscape without direct competition. This mechanism aligns well with experimental observations from surface plasmon resonance (SPR) and bio-layer interferometry (BLI) studies showing that CR3022 binding slows ACE2 association and accelerates dissociation via allosteric modulation rather than physical occlusion [57].

While the overall dynamic footprint and network topology are broadly similar across the three antibodies, minor differences emerge in the specific residues involved and their relative contributions to network integrity. EY6A shows notable involvement of residues 365, 338, 342, and 407, which are slightly more peripheral compared to the CR3022-centric core. It also interacts more strongly with positions near the ACE2 interface, such as 495, 453, 493, and 506, potentially enhancing its modulatory influence over receptor binding dynamics. COVA1-16, in contrast, engages a broader set of core residues, including 414, 418, 393, and 397, which may reflect a more distributed mode of network interaction. Its mutational sensitivity profile includes residues at the edge of the RBM, such as 406 and 493, suggesting that COVA1-16 may be more sensitive to mutations affecting the transition between open and closed RBD states. Despite these small variations, the overall architecture of the allosteric network remains highly conserved, supporting the hypothesis that group F1 antibodies operate through a common functional mechanism, albeit with fine-tuned differences in network coupling and residue-specific interactions.

Despite not overlapping with the ACE2-binding motif, all three group F1 antibodies engage residues that are topologically linked to the receptor interface through the global interaction network. Mutational profiling of allostery confirms that disruptions in this communication pathway — especially at peripheral positions — can compromise both antibody binding and ACE2 engagement, underscoring the functional importance of long-range signaling in the RBD. This supports the idea that allostery is not a secondary or marginal effect, but rather a central mechanism of neutralization for class 4 antibodies. Binding at one site (the cryptic epitope) influences function at another (ACE2 interface), demonstrating the non-local nature of immune recognition and interference. Hence, the network-based allosteric analysis reveals that group F1 class 4 antibodies leverage a modular and evolutionarily preserved interaction framework. Their neutralization strategy relies on dynamic modulation rather than direct steric blocking, reshaping the conformational ensemble of the RBD to disfavor ACE2 engagement. While this mechanism provides robustness against escape at core residues, it also introduces vulnerabilities at the periphery of the binding epitopes. Each antibody engages a distinct subset of network hubs, yet all converge on a common dynamic outcome — interference with the efficiency of ACE2 engagement through conformational and kinetic modulation. The subtle differences in residue-level network contributions suggest that each antibody may fine-tune the allosteric signal in distinct ways, possibly influencing their susceptibility to escape mutations or their efficacy in different spike conformations. These distinctions, though minor, could become relevant in therapeutic contexts, where small shifts in binding affinity or conformational preference might determine neutralization outcomes in evolving variants.

Network analysis of group F2 antibody DH1047 revealed redistribution of the residue interaction network, with increasing localization around key positions involved in receptor binding. The high centrality residues identified for DH1047 include RBD ore elements (residues 393, 480, 362, 343, 331) as well as some ACE2-proximal residues (453, 454, 456, 457, 503, 507, 508) (Figure 10A,B). This dual involvement suggests that DH1047 engages both the conserved structural scaffold and functionally relevant regions, allowing it to exert both global conformational effects and local interference with ACE2 engagement. Structural mapping further supports this hybrid mechanism, showing that the allosteric signal now propagates more directly from the cryptic site toward the ACE2-binding motif, forming a more focused communication pathway compared to group F1 antibodies (Figure 10C). However, this increased proximity to the ACE2 interface also introduces new escape vulnerabilities, particularly at T376, K378, and R408, where mutations such as T376A and R408S — observed in Omicron BA.2 — severely compromise binding. These findings highlight a trade-off between enhanced potency and increased sensitivity to antigenic drift, positioning group F2 antibodies as an intermediate stage in the mechanistic evolution of class 4 antibodies.

**Figure 10.**
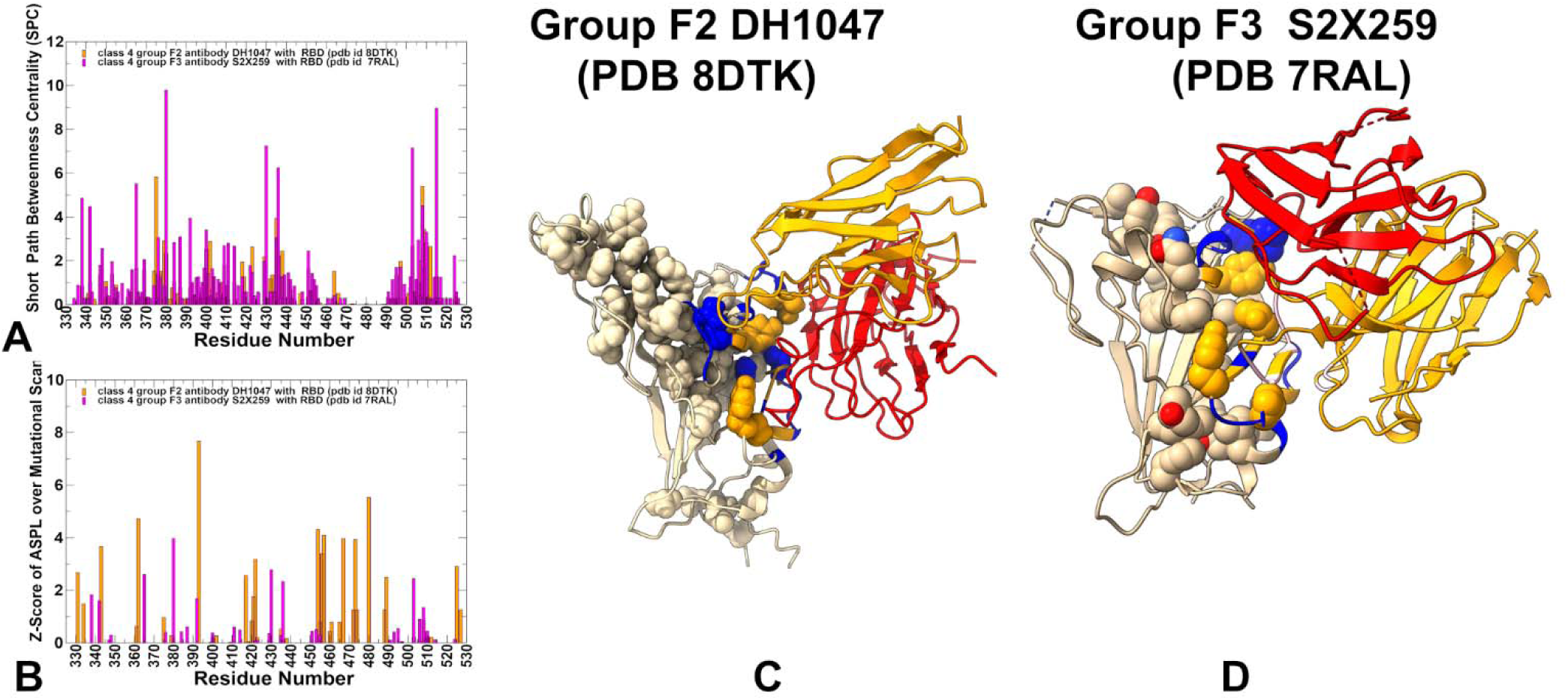
The ensemble-averaged SPC centrality (A) and the average Z-score of ASPL over mutational scan (B) for the RBD residues for class 4 group F2 antibody complex DH1047 with RBD, pdb id 8DTK (in orange filled bars) and group F3 antibody complex S2X259 with RBD, pdb id 7RAL ( in magenta filled bars). (C) Structural mapping of allosteric network sites for class 4 group F2 antibody complex DH1047 with RBD, pdb id 8DTK. (D) Structural mapping of allosteric network sites for the class 4 group F3 antibody complex S2X259 with RBD, pdb id 7RAL The heavy chain in orange ribbons, the light chain in red ribbons. The binding epitope residues are shown in blue surface, and binding hotspots are in orange and allosteric residue interaction network with high SPC and Z-score ASPL values in wheat-colored spheres.

Group F3 antibody S2X259 represents the most advanced adaptation in this trajectory, engaging the RBD through a highly localized network that overlaps extensively with the ACE2 footprint with the high centrality positions (453, 454, 457, 467, 493) that are overlap with ACE2 binding interface (Figure 10A,B). S2X259 forms more localized and direct communication pathway from the cryptic site to the receptor interface (Figure 10D), enabling dual modes of action : direct steric interference with ACE2 via overlapping contacts and allosteric reinforcement by stabilizing RBD conformations incompatible with efficient receptor engagement.

The dynamic network analysis reveals a clear evolutionary progression among class 4 antibodies, marked by increasingly focused allosteric signaling pathways that link the cryptic epitope — located deep within the RBD core — to the ACE2-binding motif on the opposite side of the domain (Figures 9,10, Table 2). This progression reflects a refinement of functional influence, transitioning from indirect allostery (F1) to hybrid mechanisms involving both conformational modulation and partial steric hindrance (F2), and finally to direct receptor competition (F3). These shifts are not only structural but also network-determined, highlighting how residue interaction patterns shape neutralization efficacy and escape vulnerability.

**Table 2.** Comparative Network Architecture for Groups F1,: F2 and F3 Antibodies: From Broad Modulation to Targeted Interference.

| **Feature** | **Group F1** | **Group F2** | **Group F3** |
| --- | --- | --- | --- |
| **Network Localization** | Broad, diffuse | Intermediate, partially localized | Highly localized |
| **ACE2 Coupling** | Weak, indirect | Moderate, hybrid | Strong, direct |
| **Key Residues** | Core β-sheet (e.g., 355–380, 431, 436) | Core + emerging ACE2 sites (T376, R408, V503) | ACE2-overlapping residues (D405, R408, G504, Y508) |
| **Escape Vulnerability** | Low | Moderate | Moderate |
| **Neutralization Mechanism** | Indirect allostery | Hybrid (dynamic + partial steric) | Direct ACE2 competition + allosteric stabilization |

## Discussion

The molecular mechanisms underlying antibody recognition of the RBD reveal a modular and progressive adaptation among class 4 antibodies — particularly across groups F1 (CR3022, EY6A, COVA1-16), group F2 (DH1047), and group F3 (S2X259) — in how they engage a deeply conserved epitope and modulate its functional consequences. While these antibodies share a common structural scaffold and conformational dependency, they exhibit divergent neutralization mechanisms, ranging from indirect allostery to direct steric competition with ACE2. This mechanistic continuum reflects an evolutionary trajectory in immune recognition, shaped by both structural constraints and selective pressures exerted during viral adaptation. Class 4 antibodies are unified by their targeting of a cryptic, hydrophobic core within the RBD, accessible only when at least two RBDs adopt the “up” conformation. This requirement introduces a conformational barrier to binding, which explains their low intrinsic potency despite broad cross-reactivity across sarbecoviruses. The structural mapping and mutational scanning data support this model, showing that escape mutations primarily affect peripheral residues, such as S383–386, T376, and R408, rather than the deeply embedded core. These findings reinforce the idea that conservation and functionally critical roles protect the central epitope from antigenic drift, while immunogenic edge regions remain mutable under selective pressure.

Dynamic simulations further illustrate how antibody binding alters the conformational landscape of the RBD, redistributing flexibility between the rigid β-sheet core and the dynamically disordered RBM loop (∼470–490). In group F1 antibodies, this redistribution enhances epitope exposure but also subtly perturbs the energetic and kinetic landscape of ACE2 engagement, consistent with experimental observations that CR3022 slows ACE2 association and accelerates dissociation, without direct physical overlap. This mechanism aligns with the hallmark of allosteric regulation, where binding at one site influences activity at another through long-range communication pathways embedded in the RBD structure. In contrast, group F2 antibody DH1047 exhibits a hybrid mode of action, extending its interaction toward the ACE2 interface and engaging key receptor-contacting residues such as T376, R408, V503, and Y508. This shift allows for partial steric hindrance, enhancing neutralization potency while still retaining significant allosteric influence. MM-GBSA and mutational profiling confirm that substitutions T376A and R408S, observed in Omicron BA.2, severely compromise DH1047 binding, highlighting the growing trade-off between breadth and escape resistance as antibodies evolve closer to the host receptor interface. Group F3 antibody S2X259 represents the most advanced stage of this progression, engaging the RBD through a highly localized network that overlaps extensively with the ACE2 footprint. This progression reflects a broader theme: neutralization strength correlates with proximity to the ACE2 interface but so does escape susceptibility. As antibodies refine their binding modes from indirect modulation to direct competition, they gain potency but lose resilience to antigenic drift. Crucially, our dynamic network modeling identifies a central “allosteric ring” embedded in the RBD core — a conserved structural motif shared across all class 4 complexes — with antibody-specific extensions propagating toward the ACE2 interface. This modular architecture supports a model in which neutralization strategies evolve via progressive refinement of peripheral network connections, rather than complete redesign of the epitope itself. This principle aligns with broader findings in protein allostery, where functional diversification occurs through modulation of network edges, while preserving the integrity of the core.

The results highlight a trade-off between breadth and potency. Group F1 antibodies (CR3022, EY6A, COVA1-16) offer broad cross-reactivity and strong resistance to escape mutations, making them ideal candidates for targeting diverse sarbecoviruses. Group F2 antibodies (DH1047) strike a functional balance, engaging both the conserved core and emerging contacts near the ACE2 interface, thereby enhancing neutralization while retaining moderate conservation. Group F3 antibodies (S2X259) achieve the highest neutralization potency by directly competing with ACE2, though at the cost of increased escape susceptibility, particularly under pressure from Omicron BA.2 variant where mutations such as T376A and R408S abolish binding.

Therapeutically, these results suggest that no single class 4 antibody is optimal in isolation. Rather, rational combinations — especially those pairing group F1 antibodies (for stability and breadth) with group F3 antibodies (for potency and direct blocking) — may offer the best balance of breadth, efficacy, and durability. Group F2 antibodies such as DH1047 could act as intermediaries, reinforcing both dynamic modulation and receptor interference without full overlapping. From a vaccine design perspective, the discovery of a structurally invariant yet dynamically influential core suggests that future immunogens should aim to stabilize open conformations of the RBD, thereby promoting recognition of cryptic epitopes that are otherwise inaccessible to the immune system. This would require innovative antigen design strategies that mimic the conformational transitions necessary for F1-class antibody engagement. Finally, the integration of mutational scanning, binding free energy decomposition, and network centrality metrics highlights the importance of considering not just direct contacts, but also network-level effects and second-order conformational changes when assessing antibody function.

## Conclusions

This study provides a comprehensive mechanistic framework for understanding how class 4 antibodies modulate the RBD conformational dynamics and neutralize the virus through indirect or direct interference with ACE2 binding. By integrating structural analysis, MD simulations, binding free energy decomposition, and network-based allosteric modeling, we uncover a progressive evolution in antibody function — from classic allostery (group F1) to hybrid mechanisms involving both dynamic modulation and partial steric hindrance (group F2), and finally to direct competitive inhibition (group F3). The agreement between DMS experiments, mutational scanning and MM-GBSA binding free energy calculations underscores the accuracy of computational methods in predicting residue-level contributions to antibody recognition. Importantly, dynamic network analysis reveals a central “allosteric ring” embedded in the RBD core, serving as a conserved communication hub across all class 4 complexes. Antibody-specific extensions from this core propagate toward the ACE2 interface, forming long-range signaling pathways that influence viral entry without direct physical overlapping. These insights reinforce the idea that neutralization need not rely on direct blocking, but can emerge through conformational redistribution and kinetic modulation, offering a non-canonical yet potent mechanism of action. Together, these observations support a modular model of RBD allostery, where neutralization strategies evolve via refinement of peripheral network connections, rather than complete redesign of the epitope itself. This evolutionary logic aligns with fundamental principles of protein allostery, reinforcing that functionally distant sites can exert meaningful influence through network-level interactions. Our findings emphasize the importance of rational antibody cocktails that combine group F1 (for breadth and stability) with group F3 (for direct receptor competition), using group F2 antibodies as intermediaries to bridge these extremes.

Together, the findings of this study support a modular model of antibody-induced allostery where neutralization strategies evolve via refinement of peripheral network connections, rather than complete redesign of the epitope itself. This evolutionary logic aligns with fundamental principles of protein allostery, reinforcing that functionally distant sites can exert meaningful influence through network-level interactions.

## Author Contributions

Conceptualization, G.V.; Methodology, M.A., V.P., B.F., G.V.; Software, M.A., V.P., B.F., G.V.; Validation, G.V.; Formal analysis, G.V., M.A., V.P.,B.F.; Investigation, G.V.; Resources, G.V., M.A., G.V. ; Data curation, M.A., G.C., G.V.; Writing— original draft preparation, G.V.; Writing—review and editing, G.V.; Visualization,. A.; G.V. Supervision G.V. Project administration, G.V.; Funding acquisition, G.V. All authors have read and agreed to the published version of the manuscript.

## Data Availability Statement

Data is fully contained within the article and Supplementary Materials. Crystal structures were obtained and downloaded from the Protein Data Bank (http://www.rcsb.org). The rendering of protein structures was done with UCSF ChimeraX package (https://www.rbvi.ucsf.edu/chimerax/) and Pymol (https://pymol.org/2/). All mutational heatmaps were produced using the developed software that is freely available at https://alshahrani.shinyapps.io/HeatMapViewerApp/.

## Acknowledgments

The authors acknowledge support from Schmid College of Science and Technology at Chapman University for providing computing resources at the Keck Center for Science and Engineering.

