## Supplemental Tables S1-S5 for "Deciphering the Mechanistic Continuum of Broadly Neutralizing Class 4 Antibodies Targeting Conserved Cryptic Epitopes of the SARS-CoV-2 Spike Protein : Operating at the Intersection of Binding, Allostery and Immune Escape"

**Table S1.** The list of the intermolecular contacts in the structure of the CR3022 complex with RBD (pdb id 6YM0).

| RBD Residue | RBD Residue Number | RBD chain | Ab Residue | Ab Residue Number | Ab chain |
|---|---|---|---|---|---|
| TYR | 369 | E | ILE | 30 | H |
| TYR | 369 | E | THR | 31 | H |
| TYR | 369 | E | GLY | 28 | H |
| TYR | 369 | E | PHE | 29 | H |
| TYR | 369 | E | TYR | 27 | H |
| ASN | 370 | E | GLY | 28 | H |
| ASN | 370 | E | TYR | 27 | H |
| SER | 371 | E | ILE | 30 | H |
| PHE | 374 | E | ILE | 30 | H |
| SER | 375 | E | TYR | 52 | H |
| SER | 375 | E | ILE | 30 | H |
| SER | 375 | E | GLY | 54 | H |
| THR | 376 | E | ILE | 30 | H |
| THR | 376 | E | TYR | 52 | H |
| THR | 376 | E | GLY | 54 | H |
| THR | 376 | E | ASP | 55 | H |
| PHE | 377 | E | THR | 31 | H |
| PHE | 377 | E | TRP | 33 | H |
| PHE | 377 | E | TYR | 52 | H |
| PHE | 377 | E | TYR | 32 | H |
| PHE | 377 | E | ILE | 30 | H |
| LYS | 378 | E | ILE | 30 | H |
| LYS | 378 | E | THR | 31 | H |
| LYS | 378 | E | TRP | 33 | H |
| LYS | 378 | E | ASP | 55 | H |
| LYS | 378 | E | GLU | 57 | H |
| LYS | 378 | E | TYR | 52 | H |
| CYS | 379 | E | THR | 31 | H |
| CYS | 379 | E | TRP | 33 | H |
| CYS | 379 | E | ILE | 102 | H |
| CYS | 379 | E | SER | 100 | H |
| CYS | 379 | E | GLY | 101 | H |
| TYR | 380 | E | TRP | 33 | H |
| TYR | 380 | E | ARG | 59 | H |
| TYR | 380 | E | THR | 104 | H |
| TYR | 380 | E | ILE | 102 | H |
| TYR | 380 | E | GLU | 57 | H |
| TYR | 380 | E | GLY | 101 | H |
| TYR | 380 | E | SER | 103 | H |

| | | | | | |
|---|---|---|---|---|---|
| GLY | 381 | E | ILE | 34 | L |
| GLY | 381 | E | TYR | 31 | L |
| GLY | 381 | E | THR | 104 | H |
| GLY | 381 | E | TYR | 38 | L |
| GLY | 381 | E | GLY | 101 | H |
| GLY | 381 | E | ILE | 102 | H |
| GLY | 381 | E | SER | 103 | H |
| GLY | 381 | E | TRP | 56 | L |
| VAL | 382 | E | GLY | 101 | H |
| VAL | 382 | E | ILE | 102 | H |
| VAL | 382 | E | SER | 103 | H |
| VAL | 382 | E | SER | 100 | H |
| VAL | 382 | E | TRP | 56 | L |
| VAL | 382 | E | ILE | 34 | L |
| VAL | 382 | E | THR | 104 | H |
| VAL | 382 | E | TYR | 38 | L |
| SER | 383 | E | GLY | 101 | H |
| SER | 383 | E | SER | 100 | H |
| SER | 383 | E | GLY | 99 | H |
| SER | 383 | E | THR | 104 | H |
| SER | 383 | E | PRO | 105 | H |
| PRO | 384 | E | SER | 100 | H |
| PRO | 384 | E | THR | 31 | H |
| PRO | 384 | E | GLY | 101 | H |
| THR | 385 | E | TYR | 32 | H |
| THR | 385 | E | THR | 31 | H |
| THR | 385 | E | SER | 100 | H |
| THR | 385 | E | GLN | 1 | H |
| THR | 385 | E | ASP | 107 | H |
| LYS | 386 | E | PRO | 105 | H |
| LYS | 386 | E | TYR | 55 | L |
| LYS | 386 | E | GLU | 61 | L |
| LYS | 386 | E | LEU | 52 | L |
| LYS | 386 | E | ASP | 107 | H |
| LYS | 386 | E | SER | 100 | H |
| ASP | 389 | E | TYR | 55 | L |
| LEU | 390 | E | TRP | 56 | L |
| PHE | 392 | E | TRP | 56 | L |
| PHE | 392 | E | ILE | 34 | L |
| ARG | 408 | E | ASP | 55 | H |
| ASP | 427 | E | TYR | 31 | L |
| ASP | 428 | E | SER | 32 | L |
| ASP | 428 | E | TYR | 31 | L |

| | | | | | |
|---|---|---|---|---|---|
| ASP | 428 | E | SER | 33 | L |
| ASP | 428 | E | TYR | 98 | L |
| PHE | 429 | E | TYR | 31 | L |
| THR | 430 | E | ILE | 34 | L |
| THR | 430 | E | TYR | 31 | L |
| THR | 430 | E | SER | 33 | L |
| THR | 430 | E | TYR | 38 | L |
| PHE | 515 | E | SER | 33 | L |
| PHE | 515 | E | ILE | 34 | L |
| GLU | 516 | E | ILE | 34 | L |
| GLU | 516 | E | SER | 33 | L |
| LEU | 517 | E | ILE | 34 | L |
| LEU | 517 | E | ASN | 35 | L |
| LEU | 517 | E | SER | 32 | L |
| LEU | 517 | E | SER | 33 | L |
| LEU | 517 | E | LYS | 36 | L |
| LEU | 518 | E | SER | 33 | L |
| HIS | 519 | E | ASN | 35 | L |

**Table S2.** The list of the intermolecular contacts in the structure of the EY6A complex with RBD (pdb id 7ZF3).

| RBD Residue | RBD Residue Number | RBD chain | Ab Residue | Ab Residue Number | Ab chain |
|---|---|---|---|---|---|
| LEU | 368 | E | TYR | 59 | H |
| LEU | 368 | E | ASN | 57 | H |
| TYR | 369 | E | ASN | 57 | H |
| TYR | 369 | E | LYS | 58 | H |
| TYR | 369 | E | SER | 56 | H |
| TYR | 369 | E | TYR | 59 | H |
| ASN | 370 | E | LYS | 58 | H |
| ASN | 370 | E | SER | 56 | H |
| ASN | 370 | E | TYR | 59 | H |
| ASN | 370 | E | ASN | 57 | H |
| ALA | 372 | E | LYS | 65 | H |
| PRO | 373 | E | GLY | 66 | H |
| PRO | 373 | E | LYS | 65 | H |
| PHE | 375 | E | LYS | 65 | H |
| THR | 376 | E | LYS | 65 | H |
| PHE | 377 | E | TYR | 59 | H |
| PHE | 377 | E | LYS | 65 | H |
| PHE | 377 | E | LEU | 95 | L |
| LYS | 378 | E | LEU | 95 | L |
| LYS | 378 | E | SER | 93 | L |
| LYS | 378 | E | ALA | 96 | L |
| LYS | 378 | E | ASP | 1 | L |
| LYS | 378 | E | ASP | 62 | H |
| CYS | 379 | E | LEU | 95 | L |
| CYS | 379 | E | SER | 93 | L |
| CYS | 379 | E | TYR | 92 | L |
| CYS | 379 | E | THR | 94 | L |
| TYR | 380 | E | SER | 93 | L |
| TYR | 380 | E | TYR | 92 | L |
| TYR | 380 | E | THR | 94 | L |
| GLY | 381 | E | SER | 93 | L |
| GLY | 381 | E | TYR | 92 | L |
| GLY | 381 | E | THR | 94 | L |
| GLY | 381 | E | TRP | 104 | H |
| GLY | 381 | E | SER | 91 | L |
| GLY | 381 | E | TYR | 32 | L |
| GLY | 381 | E | VAL | 105 | H |

| | | | | | |
|---|---|---|---|---|---|
| VAL | 382 | E | TRP | 104 | H |
| VAL | 382 | E | TYR | 32 | L |
| VAL | 382 | E | VAL | 105 | H |
| VAL | 382 | E | TYR | 92 | L |
| VAL | 382 | E | THR | 94 | L |
| SER | 383 | E | TRP | 104 | H |
| SER | 383 | E | VAL | 105 | H |
| SER | 383 | E | THR | 94 | L |
| SER | 383 | E | TYR | 106 | H |
| PRO | 384 | E | TYR | 59 | H |
| PRO | 384 | E | THR | 94 | L |
| PRO | 384 | E | TYR | 106 | H |
| PRO | 384 | E | LEU | 95 | L |
| PRO | 384 | E | ASN | 57 | H |
| THR | 385 | E | VAL | 50 | H |
| THR | 385 | E | TYR | 53 | H |
| THR | 385 | E | TYR | 106 | H |
| THR | 385 | E | SER | 52 | H |
| THR | 385 | E | TYR | 59 | H |
| THR | 385 | E | ASP | 33 | H |
| THR | 385 | E | ASN | 57 | H |
| THR | 385 | E | ILE | 51 | H |
| LYS | 386 | E | TRP | 104 | H |
| LYS | 386 | E | ASP | 33 | H |
| LYS | 386 | E | GLY | 101 | H |
| LYS | 386 | E | VAL | 105 | H |
| LYS | 386 | E | LEU | 103 | H |
| LYS | 386 | E | ASP | 99 | H |
| LYS | 386 | E | TYR | 106 | H |
| LYS | 386 | E | LYS | 102 | H |
| ASN | 388 | E | TYR | 53 | H |
| ASP | 389 | E | TYR | 53 | H |
| LEU | 390 | E | TRP | 104 | H |
| PHE | 392 | E | TRP | 104 | H |
| ALA | 411 | E | GLN | 27 | L |
| PRO | 412 | E | GLN | 27 | L |
| PRO | 412 | E | TYR | 92 | L |
| GLY | 413 | E | GLN | 27 | L |
| GLN | 414 | E | GLN | 27 | L |
| PRO | 426 | E | TYR | 92 | L |
| ASP | 427 | E | SER | 28 | L |
| ASP | 427 | E | SER | 30 | L |
| ASP | 427 | E | TYR | 92 | L |

| | | | | | |
|---|---|---|---|---|---|
| ASP | 428 | E | SER | 30 | L |
| ASP | 428 | E | TYR | 92 | L |
| PHE | 429 | E | TYR | 92 | L |
| THR | 430 | E | TYR | 92 | L |
| THR | 430 | E | TRP | 104 | H |
| LEU | 517 | E | TRP | 104 | H |

**Table S3.** The list of the intermolecular contacts in the structure of the COVA1-16 complex with RBD (pdb id 7JMW).

| RBD Residue | RBD Residue Number | RBD chain | Ab Residue | Ab Residue Number | Ab chain |
|---|---:|---|---|---:|---|
| LEU | 368 | A | ARG | 100 | H |
| TYR | 369 | A | ARG | 100 | H |
| ASN | 370 | A | ARG | 100 | H |
| SER | 371 | A | ARG | 100 | H |
| ALA | 372 | A | ARG | 100 | H |
| PHE | 374 | A | ARG | 100 | H |
| PHE | 377 | A | TYR | 100 | H |
| PHE | 377 | A | TYR | 99 | H |
| PHE | 377 | A | ARG | 100 | H |
| LYS | 378 | A | TYR | 100 | H |
| LYS | 378 | A | TYR | 99 | H |
| CYS | 379 | A | ASN | 98 | H |
| CYS | 379 | A | TYR | 100 | H |
| CYS | 379 | A | TYR | 99 | H |
| TYR | 380 | A | ARG | 97 | H |
| TYR | 380 | A | TYR | 100 | H |
| TYR | 380 | A | TYR | 99 | H |
| TYR | 380 | A | ASN | 98 | H |
| GLY | 381 | A | ASN | 98 | H |
| GLY | 381 | A | ARG | 97 | H |
| GLY | 381 | A | TYR | 100 | H |
| VAL | 382 | A | TYR | 100 | H |
| SER | 383 | A | TYR | 100 | H |
| SER | 383 | A | GLY | 100 | H |
| PRO | 384 | A | TYR | 100 | H |
| PRO | 384 | A | ARG | 100 | H |
| PRO | 384 | A | GLY | 100 | H |
| THR | 385 | A | ARG | 100 | H |
| THR | 385 | A | GLY | 100 | H |
| ARG | 408 | A | LEU | 54 | L |
| ARG | 408 | A | ASN | 53 | L |
| ARG | 408 | A | TYR | 49 | L |
| PRO | 412 | A | ARG | 97 | H |
| PRO | 412 | A | PRO | 96 | H |
| PRO | 412 | A | TYR | 32 | H |
| GLY | 413 | A | GLN | 101 | H |
| GLY | 413 | A | TYR | 32 | H |
| GLY | 413 | A | ARG | 94 | H |
| GLY | 413 | A | HIS | 102 | H |

| | | | | | |
|---|---|---|---|---|---|
| GLN | 414 | A | PRO | 96 | H |
| GLN | 414 | A | GLN | 101 | H |
| GLN | 414 | A | GLU | 55 | L |
| GLN | 414 | A | TYR | 32 | H |
| GLN | 414 | A | TYR | 49 | L |
| THR | 415 | A | THR | 56 | L |
| GLY | 416 | A | THR | 56 | L |
| PRO | 426 | A | ARG | 97 | H |
| ASP | 427 | A | TYR | 32 | H |
| ASP | 427 | A | THR | 28 | H |
| ASP | 427 | A | GLY | 26 | H |
| ASP | 427 | A | ARG | 97 | H |
| ASP | 427 | A | SER | 31 | H |
| ASP | 427 | A | TYR | 27 | H |
| ASP | 428 | A | THR | 28 | H |
| ASP | 428 | A | ARG | 97 | H |
| ASP | 428 | A | SER | 31 | H |
| PHE | 429 | A | ARG | 97 | H |
| THR | 430 | A | ARG | 97 | H |

**Table S4.** The list of the intermolecular contacts in the structure of the DH1047 complex with RBD (pdb id 8DTK).

| RBD Residue | RBD Residue Number | RBD chain | Ab Residue | Ab Residue Number | Ab chain |
|---|---:|---|---|---:|---|
| TYR | 369 | A | GLY | 100 | C |
| TYR | 369 | A | TYR | 52 | C |
| TYR | 369 | A | ASN | 56 | C |
| ASN | 370 | A | GLY | 54 | C |
| SER | 371 | A | ASN | 56 | C |
| ALA | 372 | A | ASN | 56 | C |
| ALA | 372 | A | THR | 57 | C |
| ALA | 372 | A | GLY | 55 | C |
| ALA | 372 | A | GLY | 54 | C |
| SER | 373 | A | ASN | 56 | C |
| PHE | 374 | A | ASN | 56 | C |
| SER | 375 | A | ASN | 56 | C |
| SER | 375 | A | LEU | 100 | C |
| SER | 375 | A | THR | 57 | C |
| SER | 375 | A | ASP | 100 | C |
| SER | 375 | A | ILE | 58 | C |
| THR | 376 | A | LEU | 100 | C |
| THR | 376 | A | ASP | 100 | C |
| PHE | 377 | A | ASP | 100 | C |
| PHE | 377 | A | GLY | 100 | C |
| LYS | 378 | A | TRP | 100 | C |
| LYS | 378 | A | GLY | 100 | C |
| LYS | 378 | A | ASP | 100 | C |
| CYS | 379 | A | GLY | 100 | C |
| CYS | 379 | A | TRP | 100 | C |
| TYR | 380 | A | TRP | 100 | C |
| PRO | 384 | A | GLY | 100 | C |
| GLY | 404 | A | LEU | 100 | C |
| ASP | 405 | A | GLN | 27 | B |
| ASP | 405 | A | SER | 93 | B |
| ASP | 405 | A | TYR | 92 | B |
| VAL | 407 | A | LEU | 100 | C |
| ARG | 408 | A | LEU | 100 | C |
| ARG | 408 | A | TYR | 27 | B |
| ARG | 408 | A | SER | 93 | B |
| ARG | 408 | A | TYR | 32 | B |
| ARG | 408 | A | TYR | 92 | B |

| | | | | | |
|---|---|---|---|---|---|
| ARG | 408 | A | TYR | 91 | B |
| GLN | 409 | A | TYR | 27 | B |
| GLN | 409 | A | TYR | 92 | B |
| GLY | 413 | A | SER | 27 | B |
| GLN | 414 | A | TRP | 100 | C |
| GLN | 414 | A | TYR | 27 | B |
| GLN | 414 | A | ASN | 28 | B |
| GLN | 414 | A | SER | 27 | B |
| THR | 415 | A | TYR | 27 | B |
| THR | 415 | A | SER | 27 | B |
| GLY | 416 | A | SER | 27 | B |
| ALA | 435 | A | LEU | 100 | C |
| ASN | 437 | A | ILE | 58 | C |
| PRO | 499 | A | GLN | 61 | C |
| THR | 500 | A | GLN | 61 | C |
| ASN | 501 | A | ASP | 1 | B |
| ASN | 501 | A | GLN | 61 | C |
| GLY | 502 | A | PRO | 95 | B |
| GLY | 502 | A | ASP | 1 | B |
| GLY | 502 | A | GLN | 61 | C |
| VAL | 503 | A | LEU | 94 | B |
| VAL | 503 | A | ILE | 58 | C |
| VAL | 503 | A | TYR | 59 | C |
| VAL | 503 | A | TRP | 47 | C |
| VAL | 503 | A | ASP | 1 | B |
| VAL | 503 | A | GLN | 61 | C |
| VAL | 503 | A | LEU | 100 | C |
| VAL | 503 | A | ALA | 60 | C |
| VAL | 503 | A | PRO | 95 | B |
| GLY | 504 | A | LEU | 94 | B |
| GLY | 504 | A | PRO | 95 | B |
| GLY | 504 | A | SER | 93 | B |
| GLY | 504 | A | LEU | 100 | C |
| TYR | 505 | A | ILE | 2 | B |
| TYR | 505 | A | LEU | 94 | B |
| TYR | 505 | A | GLN | 27 | B |
| TYR | 505 | A | ASP | 1 | B |
| TYR | 505 | A | SER | 93 | B |
| GLN | 506 | A | GLN | 61 | C |
| GLN | 506 | A | GLN | 64 | C |
| TYR | 508 | A | LEU | 100 | C |
| TYR | 508 | A | ILE | 58 | C |
| TYR | 508 | A | LEU | 94 | B |

**Table S5.** The list of the intermolecular contacts in the structure of the S2X259 complex with RBD (pdb id 7RAL).

| RBD Residue | RBD Residue Number | RBD chain | Ab Residue | Ab Residue Number | Ab chain |
|---|---|---|---|---|---|
| TYR | 369 | B | ILE | 52 | H |
| TYR | 369 | B | SER | 55 | H |
| TYR | 369 | B | TRP | 107 | H |
| TYR | 369 | B | MET | 54 | H |
| ASN | 370 | B | LYS | 74 | H |
| ASN | 370 | B | MET | 54 | H |
| ASN | 370 | B | SER | 55 | H |
| SER | 371 | B | SER | 55 | H |
| ALA | 372 | B | SER | 55 | H |
| ALA | 372 | B | MET | 57 | H |
| SER | 373 | B | MET | 57 | H |
| PHE | 374 | B | TRP | 107 | H |
| PHE | 374 | B | SER | 55 | H |
| PHE | 374 | B | MET | 57 | H |
| PHE | 374 | B | ILE | 52 | H |
| SER | 375 | B | ILE | 52 | H |
| SER | 375 | B | GLY | 108 | H |
| SER | 375 | B | ARG | 50 | H |
| SER | 375 | B | ASP | 109 | H |
| SER | 375 | B | TRP | 107 | H |
| THR | 376 | B | TRP | 107 | H |
| THR | 376 | B | ASP | 109 | H |
| THR | 376 | B | GLY | 108 | H |
| PHE | 377 | B | TYR | 105 | H |
| PHE | 377 | B | TRP | 107 | H |
| PHE | 377 | B | GLY | 108 | H |
| PHE | 377 | B | GLY | 106 | H |
| PHE | 377 | B | TYR | 104 | H |
| LYS | 378 | B | TYR | 104 | H |
| LYS | 378 | B | TYR | 105 | H |
| LYS | 378 | B | TRP | 107 | H |
| LYS | 378 | B | GLY | 108 | H |
| LYS | 378 | B | GLY | 106 | H |
| CYS | 379 | B | GLY | 106 | H |
| CYS | 379 | B | TYR | 104 | H |
| CYS | 379 | B | TYR | 105 | H |
| CYS | 379 | B | ASN | 103 | H |

| | | | | | |
|---|---|---|---|---|---|
| TYR | 380 | B | TYR | 104 | H |
| TYR | 380 | B | TYR | 105 | H |
| TYR | 380 | B | ASN | 103 | H |
| GLY | 381 | B | TYR | 105 | H |
| VAL | 382 | B | TYR | 105 | H |
| SER | 383 | B | TYR | 105 | H |
| SER | 383 | B | TYR | 32 | H |
| SER | 383 | B | PHE | 29 | H |
| PRO | 384 | B | TYR | 105 | H |
| PRO | 384 | B | TRP | 107 | H |
| PRO | 384 | B | TYR | 32 | H |
| PRO | 384 | B | MET | 54 | H |
| PRO | 384 | B | GLY | 106 | H |
| THR | 385 | B | MET | 54 | H |
| THR | 385 | B | TYR | 32 | H |
| THR | 385 | B | PHE | 29 | H |
| GLY | 404 | B | ASP | 109 | H |
| GLY | 404 | B | TYR | 33 | L |
| ASP | 405 | B | ASN | 26 | L |
| ASP | 405 | B | ALA | 31 | L |
| ASP | 405 | B | SER | 95 | L |
| ASP | 405 | B | TYR | 33 | L |
| GLU | 406 | B | TYR | 33 | L |
| VAL | 407 | B | ASP | 109 | H |
| VAL | 407 | B | TYR | 33 | L |
| ARG | 408 | B | ASP | 34 | L |
| ARG | 408 | B | TYR | 33 | L |
| ARG | 408 | B | ASP | 110 | H |
| ARG | 408 | B | GLY | 32 | L |
| ARG | 408 | B | ALA | 31 | L |
| ASN | 501 | B | LEU | 97 | L |
| ASN | 501 | B | SER | 98 | L |
| GLY | 502 | B | SER | 98 | L |
| GLY | 502 | B | GLY | 99 | L |
| GLY | 502 | B | SER | 95 | L |
| GLY | 502 | B | SER | 96 | L |
| GLY | 502 | B | LEU | 97 | L |
| VAL | 503 | B | PRO | 100 | L |
| VAL | 503 | B | SER | 98 | L |
| VAL | 503 | B | GLY | 99 | L |
| VAL | 503 | B | SER | 96 | L |
| VAL | 503 | B | LEU | 97 | L |
| VAL | 503 | B | SER | 95 | L |

| | | | | | |
|---|---|---|---|---|---|
| VAL | 503 | B | TYR | 93 | L |
| VAL | 503 | B | ASP | 94 | L |
| GLY | 504 | B | SER | 95 | L |
| GLY | 504 | B | SER | 96 | L |
| TYR | 505 | B | SER | 95 | L |
| GLN | 506 | B | GLY | 99 | L |
| GLN | 506 | B | SER | 98 | L |
| TYR | 508 | B | TYR | 93 | L |
| TYR | 508 | B | PRO | 100 | L |
